# Unexpected absence of a multiple-queen supergene haplotype from supercolonial populations of *Formica* ants

**DOI:** 10.1101/2024.09.15.613148

**Authors:** German Lagunas-Robles, Zul Alam, Alan Brelsford

**Affiliations:** Department of Biology, Indiana University, Bloomington, IN 47405, USA; Department of Evolution, Ecology, and Organismal Biology, University of California, Riverside, Riverside, CA 92521, USA

**Keywords:** Supergene, supercolony, polygyny, polydomy, *Formica*, ants

## Abstract

Ants exhibit many complex social organization strategies. One particularly elaborate strategy is supercoloniality, in which a colony consists of many interconnected nests (=polydomy) with many queens (=polygyny). In many species of *Formica* ants, an ancient queen number supergene determines whether a colony is monogyne (=headed by single queen) or polygyne. The presence of the rearranged *P* haplotype typically leads colonies to be polygyne. However, the presence and function of this supergene polymorphism have not been examined in supercolonial populations. Here, we use genomic data from species in the *Formica rufa* group to determine whether the *P* haplotype leads to supercoloniality. In a *Formica paralugubris* population, we find that nests are polygyne, despite the absence of the *P* haplotype in workers. We find spatial genetic ancestry patterns in nests consistent with supercolonial organization. Additionally, we find that the *P* haplotype is also absent in workers from supercolonial *Formica aquilonia*, and *Formica aquilonia* x *polyctena* hybrid populations, but is present in some *Formica polyctena* workers. We conclude that the *P* haplotype is not necessary for supercoloniality in the *Formica rufa* group, despite its longstanding association with non-supercolonial polygyny across the *Formica* genus.

## Introduction

The undeniable success of ants is in part ascribable to their variety and complexity of social organization (Wilson 1971, Hölldobler and Wilson 1990, Bourke and Franks 1995). Some colonies consist of a single nest (= monodomy), while others consist of several interconnected nests (= polydomy; Hölldobler and Wilson 1977). Colony queen number can also vary: colonies can be headed by a single queen (= monogyne) or many queens (= polygyne) (Hölldobler and Wilson 1977). Ants have evolved many social strategies multiple times (Hölldobler and Wilson 1990, Borowiec et al. 2021, Dahan and Rabeling 2022) leading to questions as to how such diverse social strategies coevolve (e.g., Favreau et al. 2018, Rubenstein et al. 2019). Recent genetic work revealed that alternative social organization strategies have evolved convergently and are typically associated with regions of reduced recombination, or supergenes (Kay et al. 2022). Supergenes allow for discrete alternative phenotypes to be inherited as single Mendelian units through tight genetic linkage (Darlington and Mather 1949, Thompson and Jiggins 2014). Theory predicts that beneficial alleles should cluster together in regions of reduced recombination, such as rearrangements, to prevent maladaptive recombinants (Kirkpatrick and Barton 2006, Yeaman 2013). Additionally, modeling work shows that social polymorphisms can result from genetic linkage between dispersal ability and social traits (Mullon et al. 2018).

In *Formica* ants, alternative social strategies have been linked to a queen number supergene (e.g., Purcell et al. 2014, Lagunas-Robles et al. 2021 Scarparo et al. 2023, De Gasperin et al. 2024). An ancient supergene (approximately 30 million years old; Purcell et al. 2021) determines whether a colony is monogyne or polygyne in many *Formica* species (Brelsford et al. 2020). Generally, workers in polygyne colonies have at least one *P* haplotype, while workers in monogyne colonies are almost exclusively homozygous for the *M* haplotype (e.g., Purcell et al. 2014, Brelsford et al. 2020, Lagunas-Robles et al. 2021, Pierce et al. 2022, Scarparo et al. 2023, but see McGuire et al. 2022). Intriguingly, *Formica truncorum* and *Formica exsecta* have the queen number supergene (Brelsford et al. 2020) but can also exhibit a social strategy known as supercoloniality (Rosengren et al. 1985, Elias et al. 2005, Seppä et al. 2012). This derived strategy has evolved independently in several ant lineages (Helanterä 2022), including *Formica* ants (Seifert 2018). Supercoloniality is an exaggerated form of polydomous, polygynous social organization with genetic connectivity and resource sharing between neighboring nests (Debout et al. 2007, Gordon and Heller 2012, Helanterä 2022). This allows for a colony to be interconnected over a broad geographic distribution. Supercolonial species exhibit a similar suite of traits classified under polygyne syndrome (Keller 1993, Keller 1995), such as dispersal by budding, local mating, and local queen recruitment. While it is important to note that not all polydomous colonies are polygynous (Hölldobler and Wilson 1977, Debout et al. 2007), it has been suggested that stable polygyne nests have the potential to become “incipient supercolonies” given favorable ecological conditions (Pedersen et al. 2006, Helanterä 2009, Huszár et al. 2014).

Supercoloniality was inferred to have multiple origins in *Formica* ants (e.g. *Formica rufa* group, *Formica exsecta* group, *Formica uralensis* group, *Formica integra* group, *Formica fusca* group; Borowiec et al. 2021). The species in the *F. rufa* group serve as an excellent model to examine the evolution of supercoloniality as the clade is socially polymorphic with several species that can form supercolonies (Seifert 2016, Seifert 2021). Of particular interest within the *F. rufa* group is the *F. lugubris* species complex (Seifert 2021). This species complex includes *Formica paralugubris* (Seifert 1996) which is characterized by its polygynous and supercolonial nature. Prior work in the Swiss Alps showed that *F. paralugubris* forms small neighboring supercolonies (Chapuisat et al. 1997, Holzer et al. 2009), with a notable lack of aggression between workers from the neighboring nests (Chapuisat et al. 2005, Holzer et al. 2006). While many large supercolonies typically exhibit near zero relatedness and show aggression to members of neighboring supercolonies (e.g. Elias et al. 2005, Pamilo et al. 2005, Pedersen et al. 2006, Thomas et al. 2006), *F. paralugubris* is tolerant of non-nestmates and does not show behavioral supercolony boundaries (Holzer et al. 2006). Despite numerous studies on supercolonies in both invasive (e.g., Giraud et al. 2002, Sunamura et al. 2009, Van Wilgenburg et al. 2010, Lenoir et al. 2016, Sorger et al. 2017) and native contexts (e.g. Elias et al. 2005, Holzer et al. 2009, Wiezik et al. 2017, Hakala et al. 2020), the genetic mechanism by which supercoloniality is determined remains elusive (Helanterä 2022).

In this study, we examine the potential link between an ancient queen number supergene and supercoloniality in three species from the *F. rufa* group. We hypothesize that if supercoloniality is an extension of polygyny, as has been shown in some species (e.g., Pedersen et al. 2006, Huszár et al. 2014), then the polygyne-associated *P* haplotype should be found in supercolonial populations of *Formica* ants. We assess the potential for a supercolonial *F. paralugubris* population by examining the spatial distribution of genetic ancestry within nests. If multiple supercolonies are present in the population, we would expect to find spatially restricted ancestry groups each spanning multiple nearby nests within a supercolony, and little admixture between supercolonies. We then assess the generality of our findings by reanalyzing published genomic data from two additional species in the *F. rufa* group (*F. aquilonia* and *F. polyctena*) from various supercolonial populations. If there is a similar genetic underpinning to supercoloniality in the *Formica rufa* group, we would expect our findings in *F. paraluguburis* to extend to *F. aquilonia* and *F. polyctena* workers from different supercolonies.

## Methods

### Nest sample collection and library preparation

In August 2018 and 2021, we collected worker ants from 41 total *F. rufa* group nests in the valley adjacent to Bosco Gurin, Ticino, Switzerland (46.3164° N, 8.4927° E). We recorded nest locations using a Garmin eTrex GPS unit.

We extracted DNA from up to 5 workers from each of 5 nests collected in 2018. The DNA was extracted with the following steps: we manually ground the head and thorax of each ant in liquid nitrogen then followed the Qiagen DNEasy insect tissue protocol for genomic DNA extraction. We eluted in 30μL of AE buffer. The samples were then prepared for ddRAD sequencing using the Brelsford et al. 2016 protocol, which implements elements proposed by Parchman et al. 2012 and Peterson et al. 2012, with restriction enzymes EcoRI and MseI. We sent the pooled library to Novogene for sequencing on a partial HiSeq X Ten lane with 150bp paired-ended reads.

We extracted DNA from workers from 37 nests collected in 2021, for up to 5 workers per colony, by manually grinding the head and thorax in liquid nitrogen, and digesting the tissue overnight at 56°C in 180 μL ATL buffer and 20 μL proteinase K. We transferred the supernatant into deep-well plates, and completed the DNA extraction with the QIAcube HT extraction robot following the manufacturer’s protocol for the QiaAmp 96 DNA kit. We eluted in 100μL of elution buffer (10 M/M Tris, pH 8.0). We prepared the genomic DNA for double-digest restriction-associated DNA (ddRAD) as described above for the 2018 colonies. We sent the pooled library to University of California San Diego’s Institute for Genomic Medicine for sequencing on a partial NovaSeq 6000 sequencing lane with 150bp paired-end reads.

### Variant calling and filtering

We demultiplexed the raw ddRAD sequence data using *process_radtags* in *Stacks* version 2.60 (Catchen et al. 2013). We merged overlapping pair-end reads with *PEAR* version 0.9.11 (Zhang et al. 2014) and then aligned the reads to a reference genome of a hybrid male between the species *Formica aquilonia* and *Formica polyctena* (Nouhaud et al. 2022a, GenBank GCA_907163055.1) with *bwa-mem2* version 2.2.1 (Li 2013), using *Samtools* version 1.16 (Li et al. 2009) to sort and merge bam files. Separately, we retrieved whole-genome sequence data from NCBI for two *Formica truncorum* males to leverage the known supergene genotypes for these samples for our analyses (Brelsford et al. 2020, Table S1). For these whole-genome samples, we used PEAR version 0.9.11 with the flag -k to retain read orientation and aligned the reads to the *Formica aquilonia* x *polyctena* reference genome (GenBank GCA_907163055.1) using *bwa-mem2* version 2.2.1. Then, for the whole-genome bam files, we used *fixmate* to fill mate coordinates and marked and removed PCR duplicates using *markdup* in *Samtools* version 1.16. We used *Samtools* version 1.16 (Li et al. 2009) to sort and merge the whole-genome bam files.

We called genomic variants for the whole-genome and ddRAD samples together with *mpileup* and *call* in *bcftools* version 1.16 (Li 2011), retaining reads with a mapping quality of at least 20. We filtered the variants to create a multi-species VCF file using the following parameters: retained bi-allelic sites (--max-alleles 2), minimum depth of 8 (--minDP 8), minor allele count of at least 2 (--mac 2), up to 25% missing data per locus (--max-missing 0.75), and removed indels (--remove-indels) in *VCFtools* version 0.1.17 (Danecek et al. 2011). Hereafter, scaffolds will be referred to as chromosomes since the reference assembly is anchored to the *Formica selysi* chromosome-level assembly (Brelsford et al. 2020, GCA_009859135.1).

### Species identification

We performed a principal component analysis (PCA) in *PLINK* version 1.90b6.25 (Purcell et al. 2007), excluding chromosome 3 (--not-chr Scaffold03) since recombination is reduced on the social supergene and would create genetic structure independent of species. We retained individuals with less than 50% missing data (--mind 0.50) as determined by *PLINK* version 1.90b6.25 (Purcell et al. 2007). This analysis resulted in three distinct clusters. We sent raw reads from three representatives of each cluster to Guillaume Lavanchy, who conducted a cluster analysis combining these samples with additional ddRAD data from seven morphologically identified *F. rufa* group species (G. Lavanchy and T. Schwander, unpublished data). The inferred species identities for our three genetic clusters were *Formica paralugubris, Formica aquilonia*, and *Formica truncorum* (Figure 1A). In total, including the whole-genome resequencing data, we had genomic data for 24 *F. aquilonia*, 155 *F. paralugubris*, and 15 *F. truncorum* (Table S1). Of the 41 nests we sampled in the single locality, we found the overwhelming majority were *F. paralugubris* (*n* = 34) and a smaller number were *F. aquilonia* (*n* = 5), and *F. truncorum* (*n* = 3).

**Figure 1.**
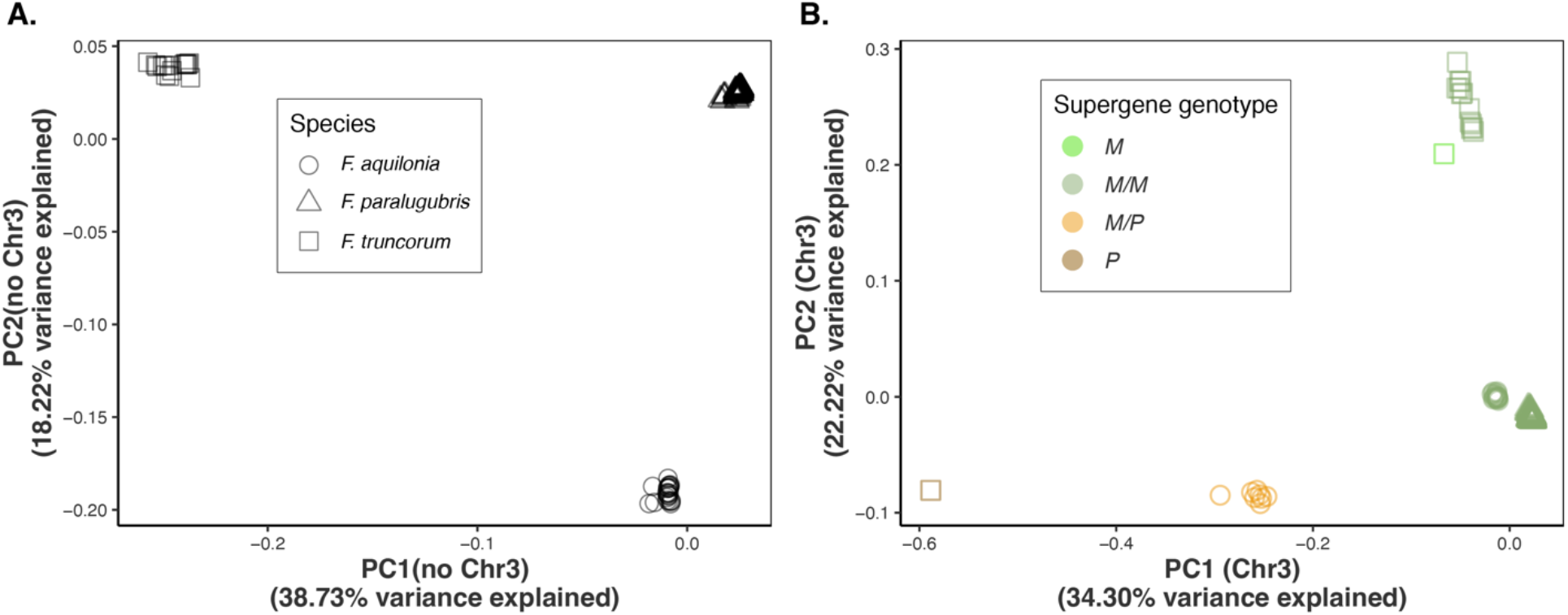
Principal component analysis (PCA) reveals that the majority of individuals in the study are *M*/*M* at the chromosome 3 supergene. Each point represents an individual. The individual’s species is represented by its shape. The individual’s haplotype (haploid males) or genotype (diploid workers) is indicated by the color. We included two whole-genome resequencing samples with known supergene genotypes for *F. truncorum* (a *M* male and a *P* male) to inform the PCA. (A) Using PCA of all SNPs excluding chromosome 3, we confirmed the presence of three species in our dataset, *F. aquilonia, F. paralugubris*, and *F. truncorum*. (B) Individuals separated into three main clusters along PC1 for chromosome 3. The single *F. truncorum* haploid *P* male was at the extreme negative end of PC1 and the homo-/hemi-zygous *M* individuals were at the extreme positive end of PC1. Notably, all the *F. paralugubris* were *M*/*M* on the supergene. The intermediate group consisted of *F. aquilonia M*/*P* workers.

### Supergene genotypes

With our filtered dataset, we generated a PCA of chromosome 3 in *PLINK* version 1.90b6.25 (Purcell et al. 2007) to identify the presence of supergene genotypes in our dataset. We only included the individuals with less than 50% missing data present that were retained in the species identification analysis. To determine the genotype heterozygosity, we calculated heterozygosity (--het) for chromosome 3 in *VCFtools* version 0.1.17 (Danecek et al. 2011). To measure the genetic distance from the reference genome, we extracted genotype calls with using “--012” in *VCFtools* version 0.1.17 (Danecek et al. 2011). In 012 format, 0 matches the reference allele, 1 is heterozygous, and 2 is homozygous for the non-reference allele. We calculated the frequency of non-reference alleles averaged across chromosome 3 for each individual (excluding missing genotypes, which are coded as -1 in this format) and converted the average to a proportion by dividing by 2.

### Estimating relatedness to infer nest social form

We filtered the raw VCF file separately to create species-specific VCF files using the following parameters: retained bi-allelic sites (--max-alleles 2), minimum depth of 8 (--minDP 8), minor allele count of 2 (--mac 2), retained loci with less than 25% missing data (--max-missing 0.75), removed indels (--remove-indels) and excluded chromosome 3 (--not-chr Scaffold03) in VCFtools *version* 0.1.17 (Danecek et al. 2011). We also excluded any individuals that had an average filtered variant depth below 8 as calculated by --depth in *VCFtools* version 0.1.17 (Danecek et al. 2011). We extracted genotype calls in 012 format in *VCFtools* version 0.1.17 (Danecek et al. 2011) for each species for 36 nests that had at least 4 sequenced workers.

We estimated intra-nest relatedness separately for each species using *COANCESTRY* version 1.0.1.10 (Wang 2011). We excluded self-comparisons from all downstream analyses. We classified nest social form using intra-nest pairwise relatedness comparisons as determined by the Wang estimator (Wang 2002) and evaluated the values against theoretical expectations. The theoretical relatedness for workers in monandrous monogyne colony is 0.75, where workers are full sisters; in a colony headed by a doubly or triply mated queen, some pairs of workers would be half sisters with expected relatedness of 0.25 (Hamilton 1964). We expect relatedness estimates to be downward biased but more precise in reduced representation datasets compared to microsatellite-based datasets (Attard et al. 2018). We called colonies with all pairwise relationships ≥ 0.55 as monogyne monandrous, colonies with bimodal distribution of pairwise relationships with at least 20% ≥ 0.55, but none ≤ 0.19 as monogyne polyandrous. We called nests polygyne if they had at least one pairwise relationship ≤ 0.19, or if the colony did not fit the monogyne criteria. We classified pairwise comparisons between workers as full-siblings (≥ 0.55), related (0.19-0.55) and unrelated (≤ 0.19).

### Fine-scale population structure in F. paralugubris

We analyzed the genetic ancestry of the *F. paralugubris* nests collected in 2021 (*n* = 32) in the population to evaluate potential presence of supercolonies. We created a bed file using the 2021 *F. paralugubris* filtered variants from the species specific VCF file, excluding chromosome 3, using the flag --make-bed in *PLINK* version 1.90b6.25 (Purcell et al. 2007). We used the bed file as input and calculated genetic ancestry (*K* = 1-10) for each worker in *ADMIXTURE* version 1.3.0 (Alexander et al. 2009). We identified the number of ancestry groups (*K*) for the population using cross-validation error scores. We then estimated the nest ancestry by averaging worker ancestry within a nest and mapped the nest according to colony GPS coordinates taken at the time of sampling.

### Analysis of the supercolonial species F. aquilonia and F. polyctena

We examined supergene variation in supercolonial populations of *F. aquilonia, F. polyctena*, and their hybrids by leveraging publicly available filtered variants in VCF format from whole-genome resequencing data (Nouhaud et al. 2022b, Portinha et al. 2022, SpecIAnt 2022). Nouhaud et al. aligned their sequence data to the same reference genome (GenBank GCA_907163055.1), so we used the filtered variants to perform subsequent analyses. We first identified the supergene genotypes on chromosome 3 by performing a principal component analysis and calculated heterozygosity on chromosome 3 using *PLINK* version 1.90b6.25 (Purcell et al. 2007) and *VCFtools* version 0.1.17 (Danecek et al. 2011), respectively. We then extracted genotype calls for chromosome 3 using the flag --012 in *VCFtools* version 0.1.17 (Danecek et al. 2011). We calculated the frequency of non-reference alleles averaged across chromosome 3 to confirm the genotypes present in *F. aquilonia, F. polyctena*, and *F. aquilonia* x *polyctena* hybrids and omitted missing genotypes as described in the “*Supergene genotypes*” methods.

### Software for figures

All plots were produced in R version 4.2.3 (R Core Team 2023) with the package *ggplot2* (Wickham 2016). We mapped nests and their respective intra-nest average ancestry in the program, 2009 QGIS version 3.30.0 RC (QGIS Development Team). We used the package *cowplot* (Wilke et al. 2024) in R version 4.2.3 to make figures 1, 2, 4, and S1 and combined the plots in figure panel 3 with Adobe Illustrator. We edited figure legends in Adobe Illustrator.

## Results

### Absence of P haplotype in a F. paralugubris population

We identified a uniform cluster for *F. paralugubris* indicating a lack of supergene polymorphism on chromosome 3 (Figure 1B). Two distinct clusters were present for *F. aquilonia* workers. The sampled *F. truncorum* workers formed one uniform cluster. We included two haploid *F. truncorum* males with known haplotypes in the PCA to identify unknown genotypes (Brelsford et al. 2020). The haploid *M* male grouped with all the sampled *F. truncorum* workers indicating they were *M*-like at the supergene (Figure 1B). The haploid *P* male did not group with any *F. truncorum* indicating we did not have any homozygous *P*-like workers (Figure 1B). The placement of the haploid males indicated that PC1 distinguished the chromosome 3 haplotypes (*M*-like vs *P*-like), while PC2 distinguished the species. One *F. aquilonia* cluster had PC1 values intermediate between the known *F. truncorum P* and *M* males on PC1 (Figure 1B). All other workers in our analysis were *M*-like homozygotes, which was further supported by heterozygosity estimates (Figures 1B, S1B). The *F. aquilonia* cluster intermediate on PC1 and with negative *F*IS suggested that these individuals were heterozygous at the supergene (Figure S1B).

To further examine supergene variation, we calculated the average frequency of non-reference alleles on chromosome 3 (Figure S1). The known *F. truncorum M* and *P* males had an average frequency of non-reference alleles across chromosome 3 of 0.152 and 0.339, respectively. The median across chromosome 3 for all *M*-like homozygous *F. truncorum* workers was 0.172. We found a similar median for all *M*-like homozygous *F. aquilonia* and *F. paralugubris* workers, 0.147 and 0.132 respectively. For the *F. aquilonia* colony with exclusively heterozygous workers, we found the median frequency of non-reference alleles to be 0.255. Through this analysis, we establish that the male sequenced for the reference genome (Nouhaud et al. 2022b) is haploid for the *M* haplotype as it is more genetically similar to the *M*-like workers than the heterozygous workers and known *P* male in our dataset. Hereafter, we will refer to the *M*-like and *P*-like haplotypes as *M* and *P*.

### Relatedness and fine-scale population structure support supercoloniality in

F. paralugubris We found support for all of the *F. paralugubris* workers belonging to polygyne nests with varying degrees of inbreeding (Figure 2A, Table S2). A history of inbreeding can increase pairwise relatedness values and complicate the inference of queen number in a nest. In nests belonging to a supercolony with frequent within-nest mating, we would expect moderately high relatedness, but not the bimodal distribution that is expected in monogyne polyandrous nests. To further evaluate supercoloniality, we estimated genetic ancestry and admixture in *F. paralugubris*. We found support for three genetic ancestry groups (*K* = 3) in the population (Figure 3, Table S3). Each genetic group included multiple nests and occupied a distinct region of our study area (Figure 3A). The spatially distinct genetic groups and consistent polygynous status suggest that this *F. paralugubris* population consists of three supercolonies.

**Figure 2.**
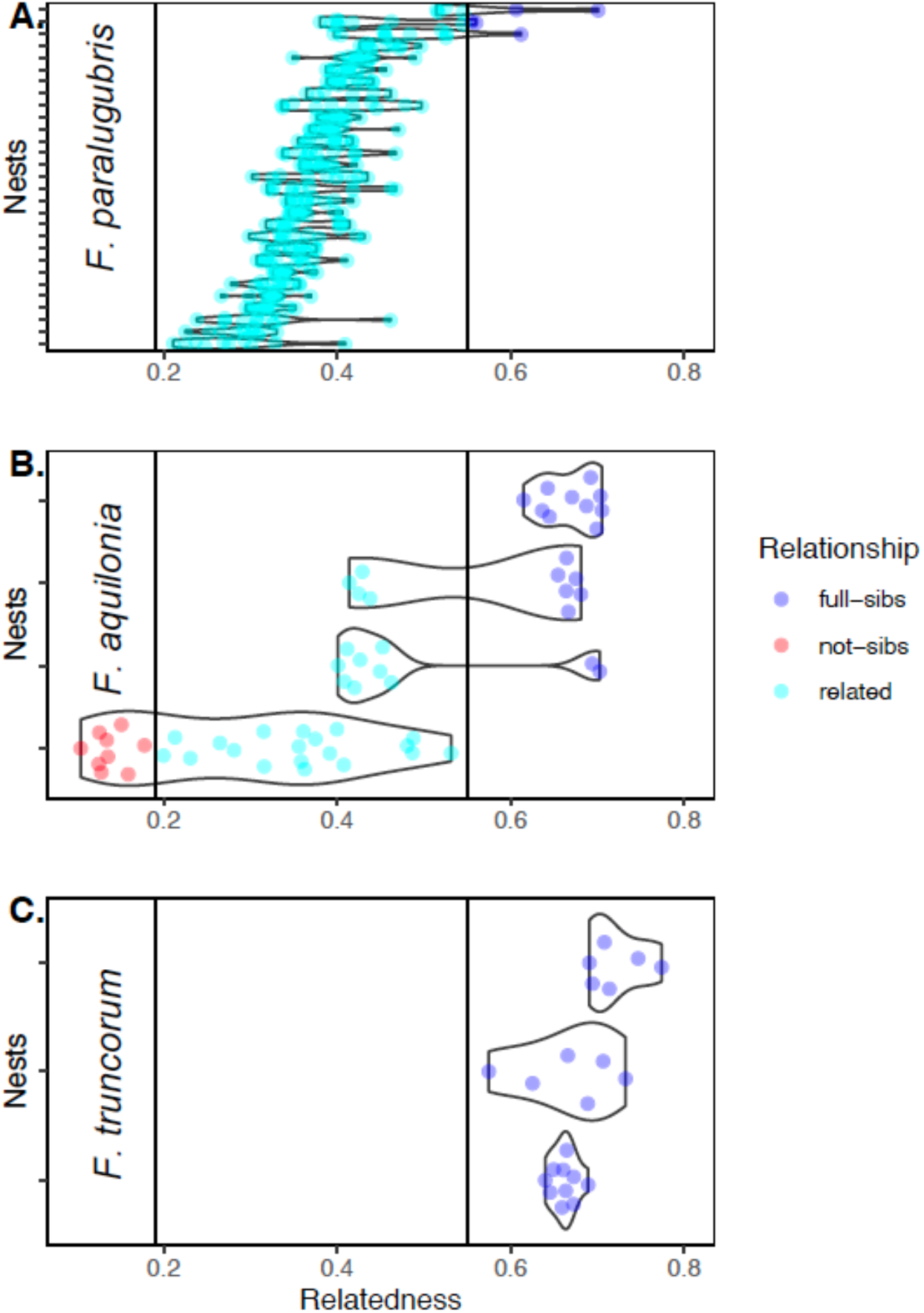
Within-nest relatedness was used to infer whether a nest was monogyne or polygyne. (A) In *F. paralugubris* (*n* = 29), we inferred all nests as polygyne. All but three nests lacked full-sibling relationships; the three nests with inferred full-sibling pairs lacked the bimodal distribution of relatedness values expected in monogyne polyandrous colonies. (B) In *F. aquilonia* (*n* = 4), three nests with the *M*/*M* genotype were inferred to be monogyne (two with polyandry) and one colony with *M*/*P* workers was inferred to be polygyne. (C) In *F. truncorum* (*n* = 3), we found all nests to be monogyne with the *M*/*M* genotype. Nests with a minimum of four sequenced workers were included in these analyses. Each point represents a comparison between two workers from the same colony. Blue points represent full-sibling relationships (pairwise R ≥ 0.55), cyan points represent related individuals (pairwise R between 0.19 and 0.55), and red points represent non-sibling relationships (pairwise R ≤ 0.19). Vertical lines at R = 0.19 and R = 0.55 show thresholds between inferred relationship types.

**Figure 3.**
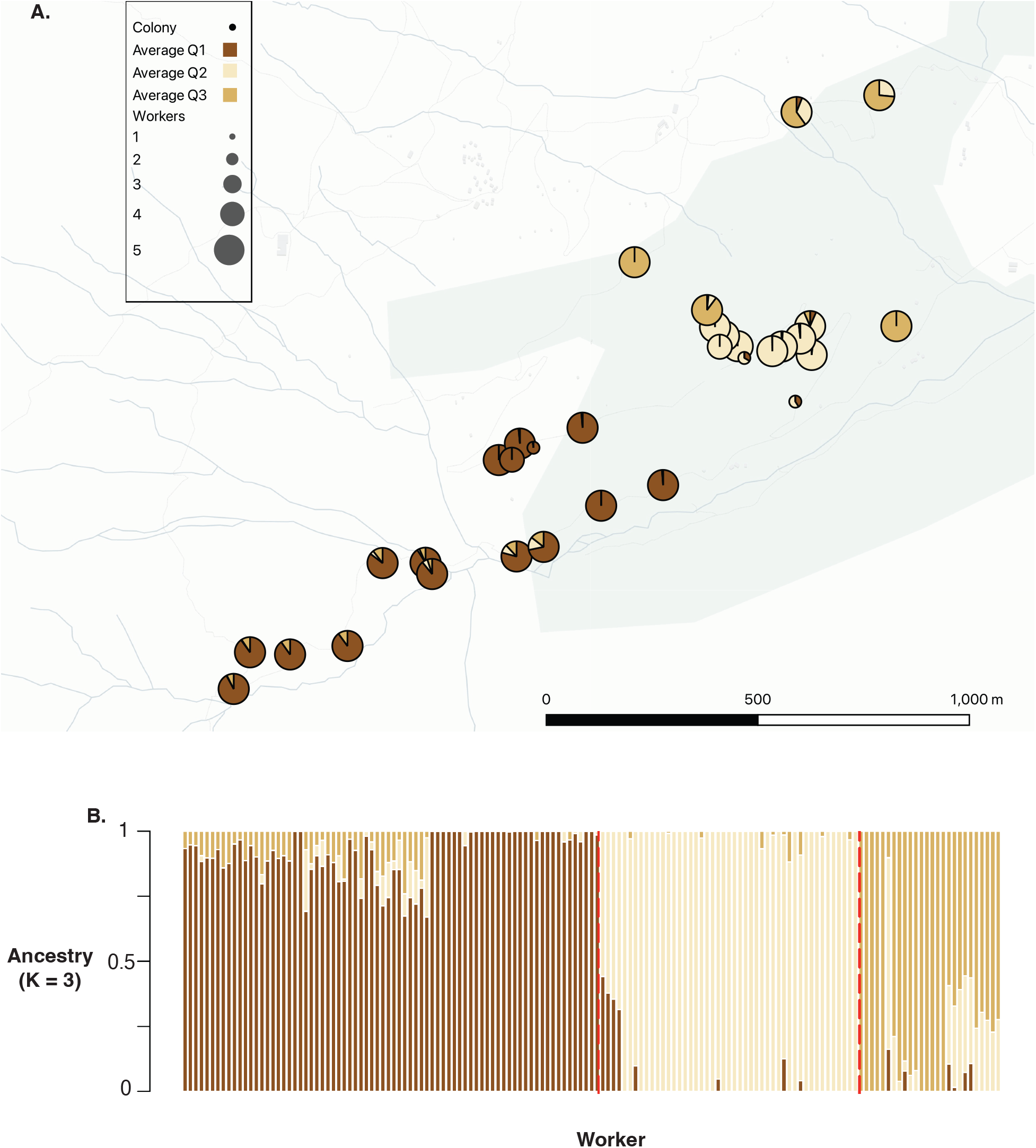
Admixture results suggest the presence of three *F. paralugubris* genetic ancestry groups. Colors represents the proportion of ancestry for each nest or individual. (A) Each nest was plotted based on its physical location. Pie charts represent the average nest ancestry. We found three genetic groups that are spatially separated. (B) ADMIXTURE results for worker ancestry (*K* = 3). Individual worker ancestries are represented by each bar. The bars are ordered by increasing latitude. The boundaries between three supercolonies are delineated by red bars.

### Supergene genotypes and social structure in F. aquilonia and F. truncorum

In our limited number of *F. aquilonia* nests (*n* = 4), nest average pairwise relatedness between workers ranged between 0.288 and 0.670 (Figure 2B, Table S4). The one nest with heterozygous *M/P* workers had the lowest mean relatedness (0.288) and a unimodal distribution of relatedness values (Figure 2B), suggesting that this colony was polygynous. We found the three *M/M* nests had intra-nest pairwise relatedness ranges between 0.401 to 0.703, 0.414 to 0.681, and 0.615 to 0.705. The pairwise relatedness in *M/M* nests suggested monogyne, monandrous social organization in one nest and monogyne, polyandrous in the other two (Figure 2B, Table S4). Intra-nest relatedness between pairs of *F. truncorum* workers ranged between 0.575 to 0.774 (Figure 2C, Table S5).

### The P haplotype is absent in supercolonial F. aquilonia and F. aquilonia x polyctena hybrid populations, but present in F. polyctena

Of 59 total individuals, only four individuals had heterozygous genotypes on chromosome 3 (Figure 4B). Upon examining supergene variation, we found that the four *F. polyctena* individuals were the only ones with a *P* haplotype, with the remaining 55 individuals being homozygous for the *M* haplotype (39 *F. aquilonia x polyctena* hybrids, 10 *F. aquilonia*, and 6 *F. polyctena*, Figure 4). We validated the genotypes for each individual by comparison to the *M* reference genome: the individuals harboring the *M/P* genotype consistently had a higher frequency of non-reference alleles compared to individuals harboring the *M*/*M* genotype (Figure 4A). These results demonstrate that the *P* haplotype is not required for supercoloniality in other *F. rufa* group species, but additional nest-level analyses are warranted to assess the effect of the *P* haplotype in *F. polyctena*.

**Figure 4.**
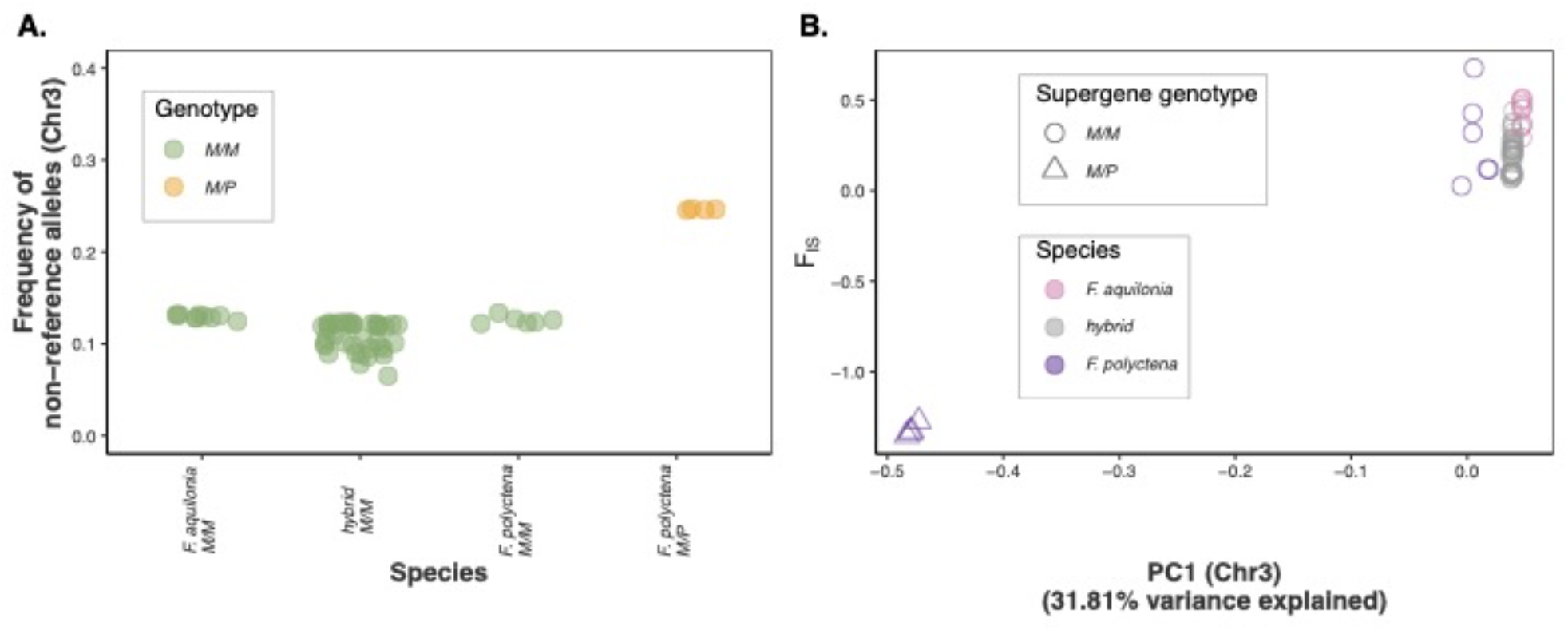
Additional species in the *F. rufa* group, *F. aquilonia* (*n* = 10), *F. polyctena* (*n* = 10), and *F. aquilonia* x *polyctena* hybrids (*n* = 59), from hybridizing supercolonial populations show that the *P* haplotype is rare in supercolonial populations. Each point represents an individual. (A) The frequency of non-reference alleles on chromosome 3 shows that four of 10 *F. polyctena* workers are substantially more divergent from the M reference genome than the rest of the workers in this dataset, suggesting that these four workers carry at least one copy of the P haplotype. (B) Chromosome 3 PC1 plotted against chromosome 3 *F*IS supported the genotype assignments as positive *F*IS values suggest the individual is homozygous on chromosome 3, while negative *F*IS values suggest the individual is heterozygous on chromosome 3. All *F. aquilonia* (*n* = 10) and *F. aquilonia* x *polyctena* hybrids (*n* = 59) were *M*/*M* at the supergene, while four of 10 *F. polyctena* were *M*/*P* at the supergene.

## Discussion

Contrary to our initial expectations, we found that the supergene haplotype associated with polygyny across the *Formica* genus is absent or rare in highly polygynous supercolonial populations of multiple species. In one population of *F. paralugubris*, nests were polygynous despite workers exclusively having the *M/M* genotype (Figure 1B, Figure 2A). This is an unexpected deviation; in other *Formica* species, colonies composed entirely of *M*/*M* genotypes are almost always monogynous (e.g., Purcell et al. 2014, Brelsford et al. 2020, Lagunas-Robles et al. 2021, Pierce et al. 2022, McGuire et al. 2022, Scarparo et al. 2023). Secondly, as expected, we find signatures of the supercolonial social organization in the population of *F. paralugubris* (Figure 2A, Figure 3). The average intra-nest ancestry suggests the presence of three supercolonies with spatial boundaries. The spatial scale of these supercolonies is consistent with studies in other populations of *F. paralugubris* (Holzer et al. 2009). Lastly, we show a lack of *P* haplotypes in additional supercolonial *Formica rufa* group species. We found this in *F. aquilonia* and *F. aquilonia* x *polyctena* hybrids, but not *F. polyctena* (Figure 4). Interestingly, in our sampling population, we observed a polygyne *F. aquilonia* nest with heterozygous *M/P* individuals (Figure 1C). This suggests that the *P* haplotype may be present in some supercolonial species, but that it is not necessary for supercolonial social organization.

### Did supercoloniality originate from polygyny?

Supercoloniality is generally thought to be an extension of polygyny (Helanterä 2009, Helanterä 2022). For example, in the common red ant *Myrmica rubra*, there are no differences in body size between individuals from polygynous and supercolony nests suggesting that morphological changes are not needed for supercoloniality (Huszár et al. 2014). However, morphological differences are observed when compared to individuals from monogyne nests. Additionally, in the Argentine ant *Linepithema humile*, differences in colony size are likely associated with local ecological conditions and not any life history changes (Pedersen et al. 2006), suggesting that native polydomous polygyne colonies (Suarez et al. 2008) expand into supercolonies given favorable environments. In contrast to these examples outside the *Formica* genus, our finding that the supergene haplotype associated with polygyny across *Formica* is absent in supercolonial populations of multiple species raises the possibility that supercoloniality can be a qualitatively different type of polygyny with a distinct origin and distinct genetic mechanism.

Some *Formica* species exhibit monogyne, polygyne, and supercolonial social strategies. In two of these species, *Formica truncorum* and *Formica exsecta*, the presence or absence of the *P* haplotype is associated with social organization in non-supercolonial populations (Brelsford et al. 2020). The same may be true of *F. aquilonia* (Figure 2B). These species present an excellent opportunity to identify the genetic basis of supercoloniality: a genome-wide association study for social organization in species exhibiting all three strategies could uncover variants that are specific to supercolonies, whether on the *M* haplotype or on other chromosomes. Identifying causal loci would provide insight into whether this life history strategy emerged once or multiple times in this set of species. In the case of a single origin in the *F. rufa* group, identifying the causal locus would also determine whether supercoloniality emerged in the common ancestor of the species group, or emerged more recently and introgressed across species boundaries.

### What is the role of the P haplotype in supercolonial Formica ants?

If the *P* haplotype is not necessary for supercoloniality, what role does the *P* haplotype play in supercolonial *Formica* species? One possibility is that some populations of facultatively supercolonial species retain the ancestral supergene-determined social polymorphism, with single-nest colonies headed by either a single queen or multiple queens depending on the presence or absence of the *P* haplotype. Another possibility, not mutually exclusive, is that the *P* haplotype is retained in supercolonial populations due to its effects on traits other than queen number. Morphological, behavioral, and population genetic evidence suggests that dispersal is typically lower for individuals from polygyne vs monogyne colonies (Keller 1993, Keller 1995 Sundström et al. 2005, Hakala et al. 2019, De Gasperin et al. 2024). If supercolonial species are subject to frequency-dependent or spatially variable selection on dispersal, this could account for the maintenance of supergene polymorphism in the absence of queen number polymorphism.

## Conclusion

We find that supercoloniality is not determined by the *P* haplotype. Despite *F. paralugubris* only having polygyne nests, all the workers were *M*/*M* at the queen number supergene. The striking deviation in *F. paralugubris* between a queen number supergene haplotype and nest phenotype contrasts with the long-standing association between the *P* haplotype and polygyny observed in previous studies of non-supercolonial *Formica* species. We observed a similar discordance in supercolonial *F. aquilonia* and *F. aquilonia x polyctena* hybrids, where all individuals lacked the *P* haplotype. Additionally, supercolonial *F. polyctena* had the *P* haplotype despite its absence in *F. aquilonia x polyctena* hybrids. Our results show that the presence of the *P* haplotype is not necessary for supercolonial organization in some species in the *F. rufa* group, but that is it nevertheless retained in socially polymorphic populations of the same species. This suggests that the evolutionary origin and genetic mechanism of supercoloniality may be qualitatively distinct from the origin and mechanism of polygyny found in single-nest colonies.

## Author contributions

Conceptualization: GLR, AB. Funding Acquisition: GLR, AB. Resources: AB. Investigation: GLR, ZA, AB. Formal Analysis: GLR, ZA. Visualization: GLR, AB. Writing – original draft: GLR. Writing – review & editing: GLR, ZA, AB. Supervision: GLR, AB.

## Acknowledgements

We sincerely thank Guillaume Lavanchy and Tanja Schwander for helping us identify the species in this study. Jessica Purcell, Giulia Scarparo, and members of the Brelsford and Purcell labs provided comments on an earlier version of this manuscript. Jessica Purcell, Tanja Schwander, and Thomas Kay helped collect samples. This material is based upon work supported by a National Science Foundation Graduate Research Fellowship (DGE-1326120) to G.L.-R., a University of California Riverside Undergraduate Education Mini-Grant to Z.A., and a National Science Foundation grant (DEB-1754834) to A.B. and Jessica Purcell. Computations were performed using the computer clusters and data storage resources of the University of California Riverside High-Performance Computing Center, which were funded by grants from National Science Foundation (MRI-2215705, MRI-1429826) and National Institutes of Health (1S10OD016290-01A1). This publication includes data generated at the University of California San Diego Institute for Genomic Medicine Genomics Center utilizing an Illumina NovaSeq 6000 that was purchased with funding from a National Institutes of Health SIG grant (S10OD026929).

## Data Availability

Worker and corresponding nest metadata are included in the supplementary material. VCF files and relatedness estimates will be deposited on Dryad. Raw sequence reads will be deposited on the Sequence Read Archive of the National Center for Biotechnology Information upon publication.

