## Supplementary material for "Unexpected absence of a multiple-queen supergene haplotype from supercolonial populations of *Formica* ants": Figure S1

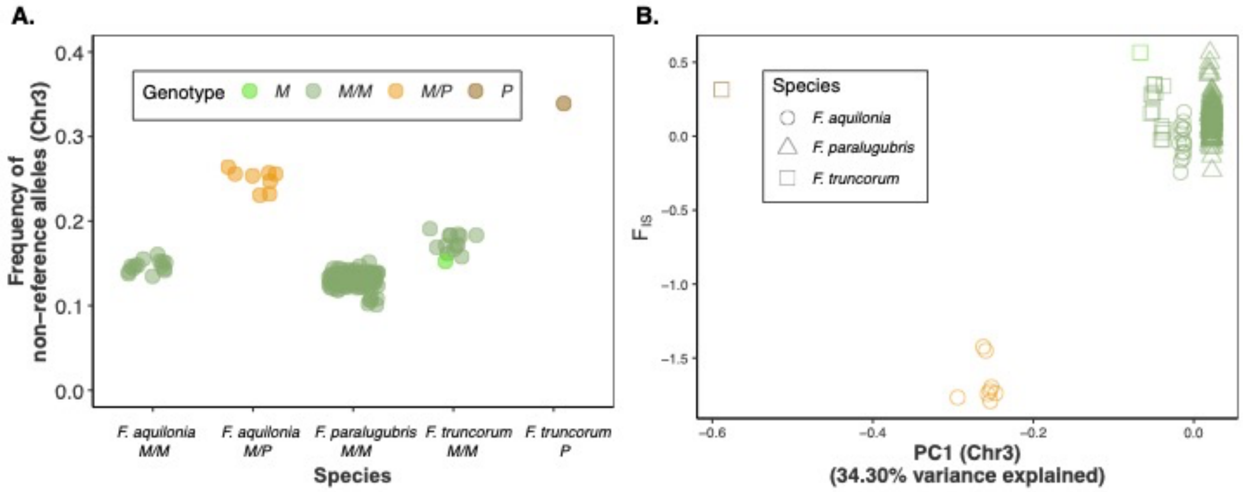


**Figure S1.** We supported the supergene assignment by using the frequency of non-reference alleles and heterozygosity on chromosome 3. (A) Eight *F. aquilonia* workers from one nest and one *F. truncorum* with the *P* haplotype had elevated divergence from the reference genome. We used two male *F. truncorum* whole-genome samples with previously identified supergene genotypes (*M* and *P* haploids): the increased distance of the *P* male from the reference suggests that the reference genome represents the *M* haplotype. We found that all *F. paralugubris* and *F. truncorum* workers had the *M*/*M* genotype, as did workers from all but *F. aquilonia* nest. (B) Chromosome 3 PC1 plotted against chromosome 3 $F_{IS}$ supported the assignment of these three clusters as *P*, *M*/*P*, and *M* homo-/hemi-zygotes (from left to right). Near zero and positive $F_{IS}$ value suggest that an individual is a homo-/hemi-zygote while a negative $F_{IS}$ value suggests that an individual is a heterozygote.

**Table S1.** Metadata for workers, inferred nest social form, chromosome 3 supergene genotype, and nest locality. Social form could not be inferred for nests with fewer than four workers. Samples included from NCBI have their corresponding accession numbers. The supergene genotypes inferred in our study from the Nouhaud et al. 2022b data are included.

| Nest | Social form | Sample | *Formica* species | Source | Chr3 | Latitude | Longitude |
|---|---|---|---|---|---|---|---|
| BG225 | Polygyne | BG225W01 | *paralugubris* | this study | *M/M* | 46.318845 | 8.48357004 |
| BG225 | Polygyne | BG225W02 | *paralugubris* | this study | *M/M* | 46.318845 | 8.48357004 |
| BG225 | Polygyne | BG225W03 | *paralugubris* | this study | *M/M* | 46.318845 | 8.48357004 |
| BG225 | Polygyne | BG225W04 | *paralugubris* | this study | *M/M* | 46.318845 | 8.48357004 |
| BG225 | Polygyne | BG225W05 | *paralugubris* | this study | *M/M* | 46.318845 | 8.48357004 |
| BG226 | Polygyne | BG226W01 | *paralugubris* | this study | *M/M* | 46.318488 | 8.481031 |
| BG226 | Polygyne | BG226W02 | *paralugubris* | this study | *M/M* | 46.318488 | 8.481031 |
| BG226 | Polygyne | BG226W03 | *paralugubris* | this study | *M/M* | 46.318488 | 8.481031 |
| BG226 | Polygyne | BG226W04 | *paralugubris* | this study | *M/M* | 46.318488 | 8.481031 |
| BG226 | Polygyne | BG226W05 | *paralugubris* | this study | *M/M* | 46.318488 | 8.481031 |
| BG227 | Monogyne | BG227W01 | *truncorum* | this study | *M/M* | 46.316162 | 8.46484403 |
| BG227 | Monogyne | BG227W02 | *truncorum* | this study | *M/M* | 46.316162 | 8.46484403 |
| BG227 | Monogyne | BG227W03 | *truncorum* | this study | *M/M* | 46.316162 | 8.46484403 |
| BG227 | Monogyne | BG227W04 | *truncorum* | this study | *M/M* | 46.316162 | 8.46484403 |
| BG227 | Monogyne | BG227W05 | *truncorum* | this study | *M/M* | 46.316162 | 8.46484403 |
| BG229-95 | Polygyne | BG229W01 | *aquilonia* | this study | *M/P* | 46.31712 | 8.47740004 |
| BG229-95 | Polygyne | BG229W02 | *aquilonia* | this study | *M/P* | 46.31712 | 8.47740004 |
| BG229-95 | Polygyne | BG229W03 | *aquilonia* | this study | *M/P* | 46.31712 | 8.47740004 |
| BG229-95 | Polygyne | BG229W04 | *aquilonia* | this study | *M/P* | 46.31712 | 8.47740004 |
| BG229-95 | Polygyne | BG229W05 | *aquilonia* | this study | *M/P* | 46.31712 | 8.47740004 |
| BG230 | <4 workers | BG230W03 | *aquilonia* | this study | *too high missing* | 46.316942 | 8.47753298 |
| BG231 | Polygyne | BG231W01 | *aquilonia* | this study | *M/M* | 46.316375 | 8.47714104 |
| BG231 | Polygyne | BG231W02 | *aquilonia* | this study | *M/M* | 46.316375 | 8.47714104 |
| BG231 | Polygyne | BG231W03 | *aquilonia* | this study | *M/M* | 46.316375 | 8.47714104 |
| BG231 | Polygyne | BG231W04 | *aquilonia* | this study | *M/M* | 46.316375 | 8.47714104 |
| BG231 | Polygyne | BG231W05 | *aquilonia* | this study | *M/M* | 46.316375 | 8.47714104 |
| BG232 | Polygyne | BG232W01 | *paralugubris* | this study | *M/M* | 46.315306 | 8.47606002 |
| BG232 | Polygyne | BG232W02 | *paralugubris* | this study | *M/M* | 46.315306 | 8.47606002 |
| BG232 | Polygyne | BG232W03 | *paralugubris* | this study | *M/M* | 46.315306 | 8.47606002 |
| BG232 | Polygyne | BG232W04 | *paralugubris* | this study | *M/M* | 46.315306 | 8.47606002 |
| BG232 | Polygyne | BG232W05 | *paralugubris* | this study | *M/M* | 46.315306 | 8.47606002 |

| | | | | | | | |
|---|---|---|---|---|---|---|---|
| BG233 | Polygyne | BG233W01 | *paralugubris* | this study | *M/M* | 46.31395 | 8.484095 |
| BG233 | Polygyne | BG233W02 | *paralugubris* | this study | *M/M* | 46.31395 | 8.484095 |
| BG233 | Polygyne | BG233W03 | *paralugubris* | this study | *M/M* | 46.31395 | 8.484095 |
| BG233 | Polygyne | BG233W04 | *paralugubris* | this study | *M/M* | 46.31395 | 8.484095 |
| BG233 | Polygyne | BG233W05 | *paralugubris* | this study | *M/M* | 46.31395 | 8.484095 |
| BG235 | Polygyne | BG235W01 | *paralugubris* | this study | *M/M* | 46.31395 | 8.48144699 |
| BG235 | Polygyne | BG235W02 | *paralugubris* | this study | *M/M* | 46.313731 | 8.48144699 |
| BG235 | Polygyne | BG235W03 | *paralugubris* | this study | *M/M* | 46.313731 | 8.48144699 |
| BG235 | Polygyne | BG235W04 | *paralugubris* | this study | *M/M* | 46.313731 | 8.48144699 |
| BG235 | Polygyne | BG235W05 | *paralugubris* | this study | *M/M* | 46.313731 | 8.48144699 |
| BG236 | Polygyne | BG236W01 | *paralugubris* | this study | *M/M* | 46.313341 | 8.48149502 |
| BG236 | Polygyne | BG236W02 | *paralugubris* | this study | *M/M* | 46.313341 | 8.48149502 |
| BG236 | Polygyne | BG236W03 | *paralugubris* | this study | *M/M* | 46.313341 | 8.48149502 |
| BG236 | Polygyne | BG236W04 | *paralugubris* | this study | *M/M* | 46.313341 | 8.48149502 |
| BG236 | Polygyne | BG236W05 | *paralugubris* | this study | *M/M* | 46.313341 | 8.48149502 |
| BG237 | Polygyne | BG237W01 | *paralugubris* | this study | *M/M* | 46.313686 | 8.481147 |
| BG237 | Polygyne | BG237W02 | *paralugubris* | this study | *M/M* | 46.313686 | 8.481147 |
| BG237 | Polygyne | BG237W03 | *paralugubris* | this study | *M/M* | 46.313686 | 8.481147 |
| BG237 | Polygyne | BG237W04 | *paralugubris* | this study | *M/M* | 46.313686 | 8.481147 |
| BG237 | Polygyne | BG237W05 | *paralugubris* | this study | *M/M* | 46.313686 | 8.481147 |
| BG238 | Polygyne | BG238W01 | *paralugubris* | this study | *M/M* | 46.313519 | 8.48058198 |
| BG238 | Polygyne | BG238W02 | *paralugubris* | this study | *M/M* | 46.313519 | 8.48058198 |
| BG238 | Polygyne | BG238W03 | *paralugubris* | this study | *M/M* | 46.313519 | 8.48058198 |
| BG238 | Polygyne | BG238W04 | *paralugubris* | this study | *M/M* | 46.313519 | 8.48058198 |
| BG238 | Polygyne | BG238W05 | *paralugubris* | this study | *M/M* | 46.313519 | 8.48058198 |
| BG239 | Polygyne | BG239W01 | *paralugubris* | this study | *M/M* | 46.313421 | 8.48028702 |
| BG239 | Polygyne | BG239W02 | *paralugubris* | this study | *M/M* | 46.313421 | 8.48028702 |
| BG239 | Polygyne | BG239W03 | *paralugubris* | this study | *M/M* | 46.313421 | 8.48028702 |
| BG239 | Polygyne | BG239W04 | *paralugubris* | this study | *M/M* | 46.313421 | 8.48028702 |
| BG239 | Polygyne | BG239W05 | *paralugubris* | this study | *M/M* | 46.313421 | 8.48028702 |
| BG240 | Polygyne | BG240W01 | *paralugubris* | this study | *M/M* | 46.313526 | 8.47922503 |
| BG240 | Polygyne | BG240W02 | *paralugubris* | this study | *M/M* | 46.313526 | 8.47922503 |
| BG240 | Polygyne | BG240W03 | *paralugubris* | this study | *M/M* | 46.313526 | 8.47922503 |
| BG240 | Polygyne | BG240W04 | *paralugubris* | this study | *M/M* | 46.313526 | 8.47922503 |
| BG240 | Polygyne | BG240W05 | *paralugubris* | this study | *M/M* | 46.313526 | 8.47922503 |
| BG241 | Polygyne | BG241W01 | *paralugubris* | this study | *M/M* | 46.31374 | 8.47880501 |
| BG241 | Polygyne | BG241W02 | *paralugubris* | this study | *M/M* | 46.31374 | 8.47880501 |
| BG241 | Polygyne | BG241W03 | *paralugubris* | this study | *M/M* | 46.31374 | 8.47880501 |
| BG241 | Polygyne | BG241W04 | *paralugubris* | this study | *M/M* | 46.31374 | 8.47880501 |

| | | | | | | | |
|---|---|---|---|---|---|---|---|
| BG241 | Polygyne | BG241W05 | *paralugubris* | this study | *M/M* | 46.31374 | 8.47880501 |
| BG242 | Polygyne | BG242W01 | *paralugubris* | this study | *M/M* | 46.313514 | 8.478674 |
| BG242 | Polygyne | BG242W03 | *paralugubris* | this study | *M/M* | 46.313514 | 8.478674 |
| BG242 | Polygyne | BG242W04 | *paralugubris* | this study | *M/M* | 46.313514 | 8.478674 |
| BG242 | Polygyne | BG242W05 | *paralugubris* | this study | *M/M* | 46.313514 | 8.478674 |
| BG243 | Polygyne | BG243W01 | *paralugubris* | this study | *M/M* | 46.313931 | 8.478529 |
| BG243 | Polygyne | BG243W02 | *paralugubris* | this study | *M/M* | 46.313931 | 8.478529 |
| BG243 | Polygyne | BG243W03 | *paralugubris* | this study | *M/M* | 46.313931 | 8.478529 |
| BG243 | Polygyne | BG243W04 | *paralugubris* | this study | *M/M* | 46.313931 | 8.478529 |
| BG243 | Polygyne | BG243W05 | *paralugubris* | this study | *M/M* | 46.313931 | 8.478529 |
| BG244 | Polygyne | BG244W01 | *paralugubris* | this study | *M/M* | 46.314291 | 8.478285 |
| BG244 | Polygyne | BG244W02 | *paralugubris* | this study | *M/M* | 46.314291 | 8.478285 |
| BG244 | Polygyne | BG244W03 | *paralugubris* | this study | *M/M* | 46.314291 | 8.478285 |
| BG244 | Polygyne | BG244W04 | *paralugubris* | this study | *M/M* | 46.314291 | 8.478285 |
| BG244 | Polygyne | BG244W05 | *paralugubris* | this study | *M/M* | 46.314291 | 8.478285 |
| BG245 | Polygyne | BG245W01 | *aquilonia* | this study | *M/M* | 46.314024 | 8.47484398 |
| BG245 | Polygyne | BG245W02 | *aquilonia* | this study | *M/M* | 46.314024 | 8.47484398 |
| BG245 | Polygyne | BG245W03 | *aquilonia* | this study | *M/M* | 46.314024 | 8.47484398 |
| BG245 | Monogyne | BG245W04 | *aquilonia* | this study | *M/M* | 46.314024 | 8.47484398 |
| BG245 | Monogyne | BG245W05 | *aquilonia* | this study | *M/M* | 46.314024 | 8.47484398 |
| BG247 | Monogyne | BG247W01 | *aquilonia* | this study | *M/M* | 46.313724 | 8.47423897 |
| BG247 | Monogyne | BG247W02 | *aquilonia* | this study | *M/M* | 46.313724 | 8.47423897 |
| BG247 | Monogyne | BG247W03 | *aquilonia* | this study | *M/M* | 46.313724 | 8.47423897 |
| BG247 | Monogyne | BG247W04 | *aquilonia* | this study | *M/M* | 46.313724 | 8.47423897 |
| BG247 | Monogyne | BG247W05 | *aquilonia* | this study | *M/M* | 46.313724 | 8.47423897 |
| BG248 | Monogyne | BG248W01 | *paralugubris* | this study | *M/M* | 46.307036 | 8.46425897 |
| BG248 | Monogyne | BG248W02 | *paralugubris* | this study | *M/M* | 46.307036 | 8.46425897 |
| BG248 | Monogyne | BG248W03 | *paralugubris* | this study | *M/M* | 46.307036 | 8.46425897 |
| BG248 | Polygyne | BG248W04 | *paralugubris* | this study | *M/M* | 46.307036 | 8.46425897 |
| BG248 | Polygyne | BG248W05 | *paralugubris* | this study | *M/M* | 46.307036 | 8.46425897 |
| BG249 | Polygyne | BG249W01 | *paralugubris* | this study | *M/M* | 46.306259 | 8.46375799 |
| BG249 | Polygyne | BG249W02 | *paralugubris* | this study | *M/M* | 46.306259 | 8.46375799 |
| BG249 | Polygyne | BG249W03 | *paralugubris* | this study | *M/M* | 46.306259 | 8.46375799 |
| BG249 | Polygyne | BG249W04 | *paralugubris* | this study | *M/M* | 46.306259 | 8.46375799 |
| BG249 | Polygyne | BG249W05 | *paralugubris* | this study | *M/M* | 46.306259 | 8.46375799 |
| BG250 | Polygyne | BG250W01 | *paralugubris* | this study | *M/M* | 46.30699 | 8.46548801 |
| BG250 | Polygyne | BG250W02 | *paralugubris* | this study | *M/M* | 46.30699 | 8.46548801 |
| BG250 | Polygyne | BG250W03 | *paralugubris* | this study | *M/M* | 46.30699 | 8.46548801 |
| BG250 | Polygyne | BG250W04 | *paralugubris* | this study | *M/M* | 46.30699 | 8.46548801 |

| | | | | | | | |
|---|---|---|---|---|---|---|---|
| BG250 | Polygyne | BG250W05 | *paralugubris* | this study | *M/M* | 46.30699 | 8.46548801 |
| BG251 | Polygyne | BG251W01 | *paralugubris* | this study | *M/M* | 46.30717 | 8.46724997 |
| BG251 | Polygyne | BG251W02 | *paralugubris* | this study | *M/M* | 46.30717 | 8.46724997 |
| BG251 | Polygyne | BG251W03 | *paralugubris* | this study | *M/M* | 46.30717 | 8.46724997 |
| BG251 | Polygyne | BG251W04 | *paralugubris* | this study | *M/M* | 46.30717 | 8.46724997 |
| BG251 | Polygyne | BG251W05 | *paralugubris* | this study | *M/M* | 46.30717 | 8.46724997 |
| BG252 | Polygyne | BG252W01 | *paralugubris* | this study | *M/M* | 46.308928 | 8.46832696 |
| BG252 | Polygyne | BG252W02 | *paralugubris* | this study | *M/M* | 46.308928 | 8.46832696 |
| BG252 | Polygyne | BG252W03 | *paralugubris* | this study | *M/M* | 46.308928 | 8.46832696 |
| BG252 | Polygyne | BG252W04 | *paralugubris* | this study | *M/M* | 46.308928 | 8.46832696 |
| BG252 | Polygyne | BG252W05 | *paralugubris* | this study | *M/M* | 46.308928 | 8.46832696 |
| BG253 | Polygyne | BG253W01 | *paralugubris* | this study | *M/M* | 46.308934 | 8.46964502 |
| BG253 | Polygyne | BG253W02 | *paralugubris* | this study | *M/M* | 46.308934 | 8.46964502 |
| BG253 | Polygyne | BG253W03 | *paralugubris* | this study | *M/M* | 46.308934 | 8.46964502 |
| BG253 | Polygyne | BG253W04 | *paralugubris* | this study | *M/M* | 46.308934 | 8.46964502 |
| BG253 | Polygyne | BG253W05 | *paralugubris* | this study | *M/M* | 46.308934 | 8.46964502 |
| BG254 | Polygyne | BG254W01 | *paralugubris* | this study | *M/M* | 46.308696 | 8.46984702 |
| BG254 | Polygyne | BG254W02 | *paralugubris* | this study | *M/M* | 46.308696 | 8.46984702 |
| BG254 | Polygyne | BG254W03 | *paralugubris* | this study | *M/M* | 46.308696 | 8.46984702 |
| BG254 | Polygyne | BG254W04 | *paralugubris* | this study | *M/M* | 46.308696 | 8.46984702 |
| BG254 | Polygyne | BG254W05 | *paralugubris* | this study | *M/M* | 46.308696 | 8.46984702 |
| BG257 | Polygyne | BG257W01 | *paralugubris* | this study | *M/M* | 46.309068 | 8.47243804 |
| BG257 | Polygyne | BG257W02 | *paralugubris* | this study | *M/M* | 46.309068 | 8.47243804 |
| BG257 | Polygyne | BG257W03 | *paralugubris* | this study | *M/M* | 46.309068 | 8.47243804 |
| BG257 | Polygyne | BG257W04 | *paralugubris* | this study | *M/M* | 46.309068 | 8.47243804 |
| BG257 | Polygyne | BG257W05 | *paralugubris* | this study | *M/M* | 46.309068 | 8.47243804 |
| BG258 | Polygyne | BG258W01 | *paralugubris* | this study | *M/M* | 46.309271 | 8.47326902 |
| BG258 | Polygyne | BG258W02 | *paralugubris* | this study | *M/M* | 46.309271 | 8.47326902 |
| BG258 | Polygyne | BG258W03 | *paralugubris* | this study | *M/M* | 46.309271 | 8.47326902 |
| BG258 | Polygyne | BG258W04 | *paralugubris* | this study | *M/M* | 46.309271 | 8.47326902 |
| BG258 | Polygyne | BG258W05 | *paralugubris* | this study | *M/M* | 46.309271 | 8.47326902 |
| BG259 | Polygyne | BG259W01 | *paralugubris* | this study | *M/M* | 46.310144 | 8.475034 |
| BG259 | Polygyne | BG259W02 | *paralugubris* | this study | *M/M* | 46.310144 | 8.475034 |
| BG259 | Polygyne | BG259W03 | *paralugubris* | this study | *M/M* | 46.310144 | 8.475034 |
| BG259 | Polygyne | BG259W04 | *paralugubris* | this study | *M/M* | 46.310144 | 8.475034 |
| BG259 | Polygyne | BG259W05 | *paralugubris* | this study | *M/M* | 46.310144 | 8.475034 |
| BG260 | Polygyne | BG260W01 | *paralugubris* | this study | *M/M* | 46.310581 | 8.47693501 |
| BG260 | Polygyne | BG260W02 | *paralugubris* | this study | *M/M* | 46.310581 | 8.47693501 |
| BG260 | Polygyne | BG260W03 | *paralugubris* | this study | *M/M* | 46.310581 | 8.47693501 |

| | | | | | | | |
|---|---|---|---|---|---|---|---|
| BG260 | Polygyne | BG260W04 | *paralugubris* | this study | *M/M* | 46.310581 | 8.47693501 |
| BG260 | Polygyne | BG260W05 | *paralugubris* | this study | *M/M* | 46.310581 | 8.47693501 |
| BG270 | Polygyne | BG270W01 | *paralugubris* | this study | *M/M* | 46.311112 | 8.47189296 |
| BG270 | Polygyne | BG270W02 | *paralugubris* | this study | *M/M* | 46.311112 | 8.47189296 |
| BG270 | Polygyne | BG270W03 | *paralugubris* | this study | *M/M* | 46.311112 | 8.47189296 |
| BG270 | Polygyne | BG270W04 | *paralugubris* | this study | *M/M* | 46.311112 | 8.47189296 |
| BG270 | Polygyne | BG270W05 | *paralugubris* | this study | *M/M* | 46.311112 | 8.47189296 |
| BG272 | Polygyne | BG272W01 | *paralugubris* | this study | *M/M* | 46.311119 | 8.47229596 |
| BG272 | Polygyne | BG272W02 | *paralugubris* | this study | *M/M* | 46.311119 | 8.47229596 |
| BG272 | Polygyne | BG272W03 | *paralugubris* | this study | *M/M* | 46.311119 | 8.47229596 |
| BG272 | Polygyne | BG272W05 | *paralugubris* | this study | *M/M* | 46.311119 | 8.47229596 |
| BG273 | Polygyne | BG273W01 | *paralugubris* | this study | *M/M* | 46.31146 | 8.47254197 |
| BG273 | Polygyne | BG273W02 | *paralugubris* | this study | *M/M* | 46.31146 | 8.47254197 |
| BG273 | Polygyne | BG273W03 | *paralugubris* | this study | *M/M* | 46.31146 | 8.47254197 |
| BG273 | Polygyne | BG273W04 | *paralugubris* | this study | *M/M* | 46.31146 | 8.47254197 |
| BG273 | Polygyne | BG273W05 | *paralugubris* | this study | *M/M* | 46.31146 | 8.47254197 |
| BG274 | Polygyne | BG274W01 | *paralugubris* | this study | *M/M* | 46.311373 | 8.47295604 |
| BG274 | Polygyne | BG274W04 | *paralugubris* | this study | *M/M* | 46.311373 | 8.47295604 |
| BG275 | Polygyne | BG275W01 | *paralugubris* | this study | *M/M* | 46.311797 | 8.47446403 |
| BG275 | Polygyne | BG275W02 | *paralugubris* | this study | *M/M* | 46.311797 | 8.47446403 |
| BG275 | Polygyne | BG275W03 | *paralugubris* | this study | *M/M* | 46.311797 | 8.47446403 |
| BG275 | Polygyne | BG275W04 | *paralugubris* | this study | *M/M* | 46.311797 | 8.47446403 |
| BG275 | Polygyne | BG275W05 | *paralugubris* | this study | *M/M* | 46.311797 | 8.47446403 |
| BG276 | <4 workers | BG276W01 | *paralugubris* | this study | *M/M* | 46.313276 | 8.47942804 |
| BG276 | <4 workers | BG276W04 | *paralugubris* | this study | *M/M* | 46.313276 | 8.47942804 |
| BG277 | <4 workers | BG277W01 | *paralugubris* | this study | *M/M* | 46.312351 | 8.48099303 |
| BG277 | <4 workers | BG277W02 | *paralugubris* | this study | *M/M* | 46.312351 | 8.48099303 |
| BG44 | Monogyne | BG44W1 | *truncorum* | this study | *M/M* | 46.314077 | 8.48766896 |
| BG44 | Monogyne | BG44W2 | *truncorum* | this study | *M/M* | 46.314077 | 8.48766896 |
| BG44 | Monogyne | BG44W3 | *truncorum* | this study | *M/M* | 46.314077 | 8.48766896 |
| BG44 | Monogyne | BG44W4 | *truncorum* | this study | *M/M* | 46.314077 | 8.48766896 |
| BG81 | Monogyne | BG81W1 | *truncorum* | this study | *M/M* | 46.312945 | 8.48624798 |
| BG81 | Monogyne | BG81W2 | *truncorum* | this study | *M/M* | 46.312945 | 8.48624798 |
| BG81 | Monogyne | BG81W3 | *truncorum* | this study | *M/M* | 46.312945 | 8.48624798 |
| BG81 | Monogyne | BG81W4 | *truncorum* | this study | *M/M* | 46.312945 | 8.48624798 |
| BG85 | <4 workers | BG85W1 | *paralugubris* | this study | *M/M* | 46.319164 | 8.48417698 |
| BG85 | <4 workers | BG85W2 | *paralugubris* | this study | *M/M* | 46.319164 | 8.48417698 |
| BG85 | <4 workers | BG85W4 | *paralugubris* | this study | *M/M* | 46.319164 | 8.48417698 |
| BG89 | <4 workers | BG89W1 | *paralugubris* | this study | *M/M* | 46.318726 | 8.48108598 |

| | | | | | | | |
|---|---|---|---|---|---|---|---|
| BG89 | <4 workers | BG89W2 | *paralugubris* | this study | *M/M* | 46.318726 | 8.48108598 |
| BG89 | <4 workers | BG89W3 | *paralugubris* | this study | *M/M* | 46.318726 | 8.48108598 |
| BG229-95 | Polygyne | BG95W1 | *aquilonia* | this study | *M/P* | 46.31712 | 8.47740004 |
| BG229-95 | Polygyne | BG95W2 | *aquilonia* | this study | *M/P* | 46.31712 | 8.47740004 |
| BG229-95 | Polygyne | BG95W3 | *aquilonia* | this study | *M/P* | 46.31712 | 8.47740004 |
| NULL | NULL | FT-MM1 | *truncorum* | Brelsford et al. 2020, SRS10116540 | *M* | NULL | NULL |
| NULL | NULL | FT-PM6 | *truncorum* | Brelsford et al. 2020, SRS10116555 | *P* | NULL | NULL |
| NULL | NULL | Att1_1w | *polyctena* | Nouhaud et al. 2022b | *M/M* | NULL | NULL |
| NULL | NULL | Bun24_10w | *aqu x polyc* hybrid | Nouhaud et al. 2022b | *M/M* | NULL | NULL |
| NULL | NULL | Bun24_11w | *aqu x polyc* hybrid | Nouhaud et al. 2022b | *M/M* | NULL | NULL |
| NULL | NULL | Bun24_12w | *aqu x polyc* hybrid | Nouhaud et al. 2022b | *M/M* | NULL | NULL |
| NULL | NULL | Bun24_8w | *aqu x polyc* hybrid | Nouhaud et al. 2022b | *M/M* | NULL | NULL |
| NULL | NULL | Bun24_9w | *aqu x polyc* hybrid | Nouhaud et al. 2022b | *M/M* | NULL | NULL |
| NULL | NULL | Bun26_4w | *aqu x polyc* hybrid | Nouhaud et al. 2022b | *M/M* | NULL | NULL |
| NULL | NULL | Bun26_5w | *aqu x polyc* hybrid | Nouhaud et al. 2022b | *M/M* | NULL | NULL |
| NULL | NULL | Bun26_6w | *aqu x polyc* hybrid | Nouhaud et al. 2022b | *M/M* | NULL | NULL |
| NULL | NULL | Bun26_7w | *aqu x polyc* hybrid | Nouhaud et al. 2022b | *M/M* | NULL | NULL |
| NULL | NULL | Bun26_8w | *aqu x polyc* hybrid | Nouhaud et al. 2022b | *M/M* | NULL | NULL |
| NULL | NULL | CAGa_1w | *polyctena* | Nouhaud et al. 2022b | *M/M* | NULL | NULL |
| NULL | NULL | CBAQ1_1w | *aquilonia* | Nouhaud et al. 2022b | *M/M* | NULL | NULL |
| NULL | NULL | CBAQ2_2w | *aquilonia* | Nouhaud et al. 2022b | *M/M* | NULL | NULL |
| NULL | NULL | CBAQ3_1w | *aquilonia* | Nouhaud et al. 2022b | *M/M* | NULL | NULL |
| NULL | NULL | CBCH1_1w | *polyctena* | Nouhaud et al. 2022b | *M/P* | NULL | NULL |
| NULL | NULL | CBCH2_2w | *polyctena* | Nouhaud et al. 2022b | *M/P* | NULL | NULL |
| NULL | NULL | CBCH3_1w | *polyctena* | Nouhaud et al. 2022b | *M//P* | NULL | NULL |
| NULL | NULL | CF14a_1w | *aquilonia* | Nouhaud et al. 2022b | *M/M* | NULL | NULL |
| NULL | NULL | CF4b_1w | *aquilonia* | Nouhaud et al. 2022b | *M/M* | NULL | NULL |

| | | | | | | | |
|---|---|---|---|---|---|---|---|
| NULL | NULL | CF8b_1w | *aquilonia* | Nouhaud et al. 2022b | *M/M* | NULL | NULL |
| NULL | NULL | FA04_4q | *aqu x polyc* hybrid | Nouhaud et al. 2022b | *M/M* | NULL | NULL |
| NULL | NULL | FA04_5q | *aqu x polyc* hybrid | Nouhaud et al. 2022b | *M/M* | NULL | NULL |
| NULL | NULL | FA04_6q | *aqu x polyc* hybrid | Nouhaud et al. 2022b | *M/M* | NULL | NULL |
| NULL | NULL | FA04_7q | *aqu x polyc* hybrid | Nouhaud et al. 2022b | *M/M* | NULL | NULL |
| NULL | NULL | FA04_8q | *aqu x polyc* hybrid | Nouhaud et al. 2022b | *M/M* | NULL | NULL |
| NULL | NULL | FA07_4q | *aqu x polyc* hybrid | Nouhaud et al. 2022b | *M/M* | NULL | NULL |
| NULL | NULL | FA07_6q | *aqu x polyc* hybrid | Nouhaud et al. 2022b | *M/M* | NULL | NULL |
| NULL | NULL | FA12_10q | *aqu x polyc* hybrid | Nouhaud et al. 2022b | *M/M* | NULL | NULL |
| NULL | NULL | FA12_11q | *aqu x polyc* hybrid | Nouhaud et al. 2022b | *M/M* | NULL | NULL |
| NULL | NULL | FA12_12q | *aqu x polyc* hybrid | Nouhaud et al. 2022b | *M/M* | NULL | NULL |
| NULL | NULL | FA12_8q | *aqu x polyc* hybrid | Nouhaud et al. 2022b | *M/M* | NULL | NULL |
| NULL | NULL | FA12_9q | *aqu x polyc* hybrid | Nouhaud et al. 2022b | *M/M* | NULL | NULL |
| NULL | NULL | FA15_4q | *aqu x polyc* hybrid | Nouhaud et al. 2022b | *M/M* | NULL | NULL |
| NULL | NULL | FA15_5q | *aqu x polyc* hybrid | Nouhaud et al. 2022b | *M/M* | NULL | NULL |
| NULL | NULL | FA16_4q | *aqu x polyc* hybrid | Nouhaud et al. 2022b | *M/M* | NULL | NULL |
| NULL | NULL | FA16_5q | *aqu x polyc* hybrid | Nouhaud et al. 2022b | *M/M* | NULL | NULL |
| NULL | NULL | FA16_6q | *aqu x polyc* hybrid | Nouhaud et al. 2022b | *M/M* | NULL | NULL |
| NULL | NULL | FAu14_2q | *aqu x polyc* hybrid | Nouhaud et al. 2022b | *M/M* | NULL | NULL |
| NULL | NULL | FAu14_5q | *aqu x polyc* hybrid | Nouhaud et al. 2022b | *M/M* | NULL | NULL |
| NULL | NULL | Fis2_1w | *polyctena* | Nouhaud et al. 2022b | *M/P* | NULL | NULL |
| NULL | NULL | Jar6_1w | *polyctena* | Nouhaud et al. 2022b | *M/M* | NULL | NULL |
| NULL | NULL | Lai_1w | *aquilonia* | Nouhaud et al. 2022b | *M/M* | NULL | NULL |
| NULL | NULL | Lai_2w | *aquilonia* | Nouhaud et al. 2022b | *M/M* | NULL | NULL |
| NULL | NULL | Loa_1w | *aquilonia* | Nouhaud et al. 2022b | *M/M* | NULL | NULL |
| NULL | NULL | Lok3_1w | *polyctena* | Nouhaud et al. 2022b | *M/M* | NULL | NULL |
| NULL | NULL | Mar1_15q | *aqu x polyc* hybrid | Nouhaud et al. 2022b | *M/M* | NULL | NULL |

| | | | | | | | |
|---|---|---|---|---|---|---|---|
| NULL | NULL | Mar1_16q | *aqu x polyc* hybrid | Nouhaud et al. 2022b | *M/M* | NULL | NULL |
| NULL | NULL | Mar1_6q | *aqu x polyc* hybrid | Nouhaud et al. 2022b | *M/M* | NULL | NULL |
| NULL | NULL | Mar1_7q | *aqu x polyc* hybrid | Nouhaud et al. 2022b | *M/M* | NULL | NULL |
| NULL | NULL | Mar1_8q | *aqu x polyc* hybrid | Nouhaud et al. 2022b | *M/M* | NULL | NULL |
| NULL | NULL | Mar2_10q | *aqu x polyc* hybrid | Nouhaud et al. 2022b | *M/M* | NULL | NULL |
| NULL | NULL | Mar2_6q | *aqu x polyc* hybrid | Nouhaud et al. 2022b | *M/M* | NULL | NULL |
| NULL | NULL | Mar2_7q | *aqu x polyc* hybrid | Nouhaud et al. 2022b | *M/M* | NULL | NULL |
| NULL | NULL | Mar2_8q | *aqu x polyc* hybrid | Nouhaud et al. 2022b | *M/M* | NULL | NULL |
| NULL | NULL | Mar2_9q | *aqu x polyc* hybrid | Nouhaud et al. 2022b | *M/M* | NULL | NULL |
| NULL | NULL | NAZa_1w | *polyctena* | Nouhaud et al. 2022b | *M/M* | NULL | NULL |
| NULL | NULL | Pus2_1w | *apquilonia* | Nouhaud et al. 2022b | *M/M* | NULL | NULL |
| NULL | NULL | VDa_1w | *polyctena* | Nouhaud et al. 2022b | *M/M* | NULL | NULL |

**Table S2.** Intra-nest relatedness as estimated by *COANCESTRY* using the Wang estimator in *F. paralugubris*. These results are visualized in the main text in Figure 2A.

| **Ind1** | **Ind2** | **Wang estimator** | **Parentage** | **Nest social form** | **Supercolony** |
|---|---|---|---|---|---|
| BG258-W03 | BG258-W05 | 0.2366 | related | polygyne | 1 |
| BG258-W02 | BG258-W05 | 0.2931 | related | polygyne | 1 |
| BG258-W03 | BG258-W04 | 0.2689 | related | polygyne | 1 |
| BG258-W04 | BG258-W05 | 0.2251 | related | polygyne | 1 |
| BG258-W01 | BG258-W03 | 0.2111 | related | polygyne | 1 |
| BG258-W02 | BG258-W04 | 0.2751 | related | polygyne | 1 |
| BG258-W01 | BG258-W05 | 0.3008 | related | polygyne | 1 |
| BG258-W01 | BG258-W02 | 0.2476 | related | polygyne | 1 |
| BG258-W02 | BG258-W03 | 0.3285 | related | polygyne | 1 |
| BG258-W01 | BG258-W04 | 0.4092 | related | polygyne | 1 |
| BG257-W01 | BG257-W05 | 0.2658 | related | polygyne | 1 |
| BG257-W02 | BG257-W03 | 0.3304 | related | polygyne | 1 |
| BG257-W01 | BG257-W02 | 0.3126 | related | polygyne | 1 |
| BG257-W01 | BG257-W04 | 0.2251 | related | polygyne | 1 |
| BG257-W02 | BG257-W04 | 0.3003 | related | polygyne | 1 |
| BG257-W01 | BG257-W03 | 0.2924 | related | polygyne | 1 |
| BG257-W03 | BG257-W05 | 0.3031 | related | polygyne | 1 |

| | | | | | |
|---|---|---|---|---|---|
| BG257-W04 | BG257-W05 | 0.2523 | related | polygyne | 1 |
| BG257-W02 | BG257-W05 | 0.3088 | related | polygyne | 1 |
| BG257-W03 | BG257-W04 | 0.2833 | related | polygyne | 1 |
| BG254-W02 | BG254-W03 | 0.3074 | related | polygyne | 1 |
| BG254-W01 | BG254-W05 | 0.3224 | related | polygyne | 1 |
| BG254-W01 | BG254-W04 | 0.2705 | related | polygyne | 1 |
| BG254-W02 | BG254-W05 | 0.4615 | related | polygyne | 1 |
| BG254-W02 | BG254-W04 | 0.2913 | related | polygyne | 1 |
| BG254-W01 | BG254-W02 | 0.3264 | related | polygyne | 1 |
| BG254-W03 | BG254-W05 | 0.3363 | related | polygyne | 1 |
| BG254-W04 | BG254-W05 | 0.2705 | related | polygyne | 1 |
| BG254-W03 | BG254-W04 | 0.2383 | related | polygyne | 1 |
| BG254-W01 | BG254-W03 | 0.3071 | related | polygyne | 1 |
| BG250-W03 | BG250-W04 | 0.3209 | related | polygyne | 1 |
| BG250-W01 | BG250-W02 | 0.2997 | related | polygyne | 1 |
| BG250-W04 | BG250-W05 | 0.3094 | related | polygyne | 1 |
| BG250-W02 | BG250-W05 | 0.3443 | related | polygyne | 1 |
| BG250-W01 | BG250-W05 | 0.3521 | related | polygyne | 1 |
| BG250-W03 | BG250-W05 | 0.3183 | related | polygyne | 1 |
| BG250-W01 | BG250-W03 | 0.3277 | related | polygyne | 1 |
| BG250-W01 | BG250-W04 | 0.3183 | related | polygyne | 1 |
| BG250-W02 | BG250-W04 | 0.3053 | related | polygyne | 1 |
| BG250-W02 | BG250-W03 | 0.2946 | related | polygyne | 1 |
| BG273-W03 | BG273-W04 | 0.3268 | related | polygyne | 1 |
| BG273-W04 | BG273-W05 | 0.3328 | related | polygyne | 1 |
| BG273-W01 | BG273-W03 | 0.3306 | related | polygyne | 1 |
| BG273-W02 | BG273-W03 | 0.3178 | related | polygyne | 1 |
| BG273-W01 | BG273-W02 | 0.3288 | related | polygyne | 1 |
| BG273-W01 | BG273-W04 | 0.2663 | related | polygyne | 1 |
| BG273-W02 | BG273-W05 | 0.3697 | related | polygyne | 1 |
| BG273-W03 | BG273-W05 | 0.3217 | related | polygyne | 1 |
| BG273-W01 | BG273-W05 | 0.2984 | related | polygyne | 1 |
| BG273-W02 | BG273-W04 | 0.32 | related | polygyne | 1 |
| BG275-W01 | BG275-W03 | 0.3054 | related | polygyne | 1 |
| BG275-W04 | BG275-W05 | 0.3386 | related | polygyne | 1 |
| BG275-W02 | BG275-W04 | 0.3151 | related | polygyne | 1 |
| BG275-W03 | BG275-W04 | 0.3565 | related | polygyne | 1 |
| BG275-W01 | BG275-W04 | 0.3121 | related | polygyne | 1 |
| BG275-W01 | BG275-W05 | 0.3271 | related | polygyne | 1 |

| | | | | | |
|---|---|---|---|---|---|
| BG275-W01 | BG275-W02 | 0.3444 | related | polygyne | 1 |
| BG275-W02 | BG275-W05 | 0.3568 | related | polygyne | 1 |
| BG275-W03 | BG275-W05 | 0.3325 | related | polygyne | 1 |
| BG275-W02 | BG275-W03 | 0.2778 | related | polygyne | 1 |
| BG249-W01 | BG249-W05 | 0.3697 | related | polygyne | 1 |
| BG249-W01 | BG249-W02 | 0.3298 | related | polygyne | 1 |
| BG249-W02 | BG249-W05 | 0.3394 | related | polygyne | 1 |
| BG249-W03 | BG249-W05 | 0.346 | related | polygyne | 1 |
| BG249-W01 | BG249-W03 | 0.3386 | related | polygyne | 1 |
| BG249-W03 | BG249-W04 | 0.3417 | related | polygyne | 1 |
| BG249-W04 | BG249-W05 | 0.3258 | related | polygyne | 1 |
| BG249-W01 | BG249-W04 | 0.3766 | related | polygyne | 1 |
| BG249-W02 | BG249-W03 | 0.3281 | related | polygyne | 1 |
| BG249-W02 | BG249-W04 | 0.3357 | related | polygyne | 1 |
| BG252-W03 | BG252-W05 | 0.4114 | related | polygyne | 1 |
| BG252-W03 | BG252-W04 | 0.3534 | related | polygyne | 1 |
| BG252-W01 | BG252-W02 | 0.3235 | related | polygyne | 1 |
| BG252-W01 | BG252-W05 | 0.3137 | related | polygyne | 1 |
| BG252-W04 | BG252-W05 | 0.3606 | related | polygyne | 1 |
| BG252-W02 | BG252-W04 | 0.3296 | related | polygyne | 1 |
| BG252-W01 | BG252-W03 | 0.3185 | related | polygyne | 1 |
| BG252-W01 | BG252-W04 | 0.3075 | related | polygyne | 1 |
| BG252-W02 | BG252-W03 | 0.3719 | related | polygyne | 1 |
| BG252-W02 | BG252-W05 | 0.3562 | related | polygyne | 1 |
| BG244-W01 | BG244-W02 | 0.3604 | related | polygyne | 2 |
| BG244-W02 | BG244-W04 | 0.3743 | related | polygyne | 2 |
| BG244-W04 | BG244-W05 | 0.3248 | related | polygyne | 2 |
| BG244-W02 | BG244-W03 | 0.3763 | related | polygyne | 2 |
| BG244-W03 | BG244-W05 | 0.3463 | related | polygyne | 2 |
| BG244-W01 | BG244-W05 | 0.3323 | related | polygyne | 2 |
| BG244-W01 | BG244-W04 | 0.3577 | related | polygyne | 2 |
| BG244-W01 | BG244-W03 | 0.3458 | related | polygyne | 2 |
| BG244-W02 | BG244-W05 | 0.3184 | related | polygyne | 2 |
| BG244-W03 | BG244-W04 | 0.3625 | related | polygyne | 2 |
| BG253-W02 | BG253-W05 | 0.3352 | related | polygyne | 1 |
| BG253-W03 | BG253-W05 | 0.3335 | related | polygyne | 1 |
| BG253-W01 | BG253-W02 | 0.3511 | related | polygyne | 1 |
| BG253-W02 | BG253-W03 | 0.3133 | related | polygyne | 1 |
| BG253-W03 | BG253-W04 | 0.2983 | related | polygyne | 1 |

| | | | | | |
|---|---|---|---|---|---|
| BG253-W01 | BG253-W04 | 0.3794 | related | polygyne | 1 |
| BG253-W01 | BG253-W03 | 0.4225 | related | polygyne | 1 |
| BG253-W04 | BG253-W05 | 0.3664 | related | polygyne | 1 |
| BG253-W02 | BG253-W04 | 0.4313 | related | polygyne | 1 |
| BG253-W01 | BG253-W05 | 0.3383 | related | polygyne | 1 |
| BG251-W03 | BG251-W05 | 0.3429 | related | polygyne | 1 |
| BG251-W03 | BG251-W04 | 0.3483 | related | polygyne | 1 |
| BG251-W04 | BG251-W05 | 0.3392 | related | polygyne | 1 |
| BG251-W02 | BG251-W05 | 0.3296 | related | polygyne | 1 |
| BG251-W02 | BG251-W04 | 0.318 | related | polygyne | 1 |
| BG251-W02 | BG251-W03 | 0.3364 | related | polygyne | 1 |
| BG251-W01 | BG251-W05 | 0.4143 | related | polygyne | 1 |
| BG251-W01 | BG251-W03 | 0.4049 | related | polygyne | 1 |
| BG251-W01 | BG251-W02 | 0.4023 | related | polygyne | 1 |
| BG251-W01 | BG251-W04 | 0.4018 | related | polygyne | 1 |
| BG235-W01 | BG235-W05 | 0.3627 | related | polygyne | 2 |
| BG235-W01 | BG235-W02 | 0.3557 | related | polygyne | 2 |
| BG235-W03 | BG235-W04 | 0.3556 | related | polygyne | 2 |
| BG235-W04 | BG235-W05 | 0.3971 | related | polygyne | 2 |
| BG235-W02 | BG235-W05 | 0.3576 | related | polygyne | 2 |
| BG235-W03 | BG235-W05 | 0.3483 | related | polygyne | 2 |
| BG235-W01 | BG235-W03 | 0.4056 | related | polygyne | 2 |
| BG235-W02 | BG235-W04 | 0.3658 | related | polygyne | 2 |
| BG235-W02 | BG235-W03 | 0.3452 | related | polygyne | 2 |
| BG235-W01 | BG235-W04 | 0.3443 | related | polygyne | 2 |
| BG259-W02 | BG259-W03 | 0.3441 | related | polygyne | 1 |
| BG259-W03 | BG259-W04 | 0.3633 | related | polygyne | 1 |
| BG259-W03 | BG259-W05 | 0.3409 | related | polygyne | 1 |
| BG259-W04 | BG259-W05 | 0.3528 | related | polygyne | 1 |
| BG259-W01 | BG259-W04 | 0.3862 | related | polygyne | 1 |
| BG259-W01 | BG259-W02 | 0.4186 | related | polygyne | 1 |
| BG259-W02 | BG259-W05 | 0.3518 | related | polygyne | 1 |
| BG259-W01 | BG259-W05 | 0.3967 | related | polygyne | 1 |
| BG259-W02 | BG259-W04 | 0.3655 | related | polygyne | 1 |
| BG259-W01 | BG259-W03 | 0.3719 | related | polygyne | 1 |
| BG248-W01 | BG248-W04 | 0.4182 | related | polygyne | 1 |
| BG248-W02 | BG248-W05 | 0.4678 | related | polygyne | 1 |
| BG248-W02 | BG248-W03 | 0.3679 | related | polygyne | 1 |
| BG248-W04 | BG248-W05 | 0.462 | related | polygyne | 1 |

| | | | | | |
|---|---|---|---|---|---|
| BG248-W03 | BG248-W05 | 0.3584 | related | polygyne | 1 |
| BG248-W01 | BG248-W03 | 0.3486 | related | polygyne | 1 |
| BG248-W02 | BG248-W04 | 0.3279 | related | polygyne | 1 |
| BG248-W03 | BG248-W04 | 0.3194 | related | polygyne | 1 |
| BG248-W01 | BG248-W02 | 0.3468 | related | polygyne | 1 |
| BG248-W01 | BG248-W05 | 0.325 | related | polygyne | 1 |
| BG260-W01 | BG260-W05 | 0.4352 | related | polygyne | 1 |
| BG260-W01 | BG260-W03 | 0.4184 | related | polygyne | 1 |
| BG260-W02 | BG260-W04 | 0.3339 | related | polygyne | 1 |
| BG260-W01 | BG260-W04 | 0.3021 | related | polygyne | 1 |
| BG260-W01 | BG260-W02 | 0.4066 | related | polygyne | 1 |
| BG260-W03 | BG260-W05 | 0.3991 | related | polygyne | 1 |
| BG260-W02 | BG260-W03 | 0.3885 | related | polygyne | 1 |
| BG260-W02 | BG260-W05 | 0.4242 | related | polygyne | 1 |
| BG260-W03 | BG260-W04 | 0.3664 | related | polygyne | 1 |
| BG260-W04 | BG260-W05 | 0.3252 | related | polygyne | 1 |
| BG239-W01 | BG239-W05 | 0.3772 | related | polygyne | 2 |
| BG239-W01 | BG239-W02 | 0.3809 | related | polygyne | 2 |
| BG239-W03 | BG239-W05 | 0.3822 | related | polygyne | 2 |
| BG239-W02 | BG239-W03 | 0.4225 | related | polygyne | 2 |
| BG239-W01 | BG239-W03 | 0.3572 | related | polygyne | 2 |
| BG239-W01 | BG239-W04 | 0.3624 | related | polygyne | 2 |
| BG239-W02 | BG239-W04 | 0.3947 | related | polygyne | 2 |
| BG239-W03 | BG239-W04 | 0.3674 | related | polygyne | 2 |
| BG239-W04 | BG239-W05 | 0.362 | related | polygyne | 2 |
| BG239-W02 | BG239-W05 | 0.3947 | related | polygyne | 2 |
| BG236-W02 | BG236-W05 | 0.3761 | related | polygyne | 2 |
| BG236-W03 | BG236-W04 | 0.3368 | related | polygyne | 2 |
| BG236-W03 | BG236-W05 | 0.3687 | related | polygyne | 2 |
| BG236-W04 | BG236-W05 | 0.3567 | related | polygyne | 2 |
| BG236-W01 | BG236-W03 | 0.3412 | related | polygyne | 2 |
| BG236-W01 | BG236-W02 | 0.4213 | related | polygyne | 2 |
| BG236-W02 | BG236-W03 | 0.4676 | related | polygyne | 2 |
| BG236-W01 | BG236-W05 | 0.3644 | related | polygyne | 2 |
| BG236-W01 | BG236-W04 | 0.4199 | related | polygyne | 2 |
| BG236-W02 | BG236-W04 | 0.3959 | related | polygyne | 2 |
| BG238-W01 | BG238-W05 | 0.3948 | related | polygyne | 2 |
| BG238-W03 | BG238-W05 | 0.4177 | related | polygyne | 2 |
| BG238-W01 | BG238-W04 | 0.3546 | related | polygyne | 2 |

| | | | | | |
|---|---|---|---|---|---|
| BG238-W02 | BG238-W05 | 0.4046 | related | polygyne | 2 |
| BG238-W02 | BG238-W03 | 0.4169 | related | polygyne | 2 |
| BG238-W01 | BG238-W02 | 0.3975 | related | polygyne | 2 |
| BG238-W02 | BG238-W04 | 0.3674 | related | polygyne | 2 |
| BG238-W01 | BG238-W03 | 0.3632 | related | polygyne | 2 |
| BG238-W03 | BG238-W04 | 0.3974 | related | polygyne | 2 |
| BG238-W04 | BG238-W05 | 0.3806 | related | polygyne | 2 |
| BG240-W01 | BG240-W03 | 0.382 | related | polygyne | 2 |
| BG240-W01 | BG240-W04 | 0.4058 | related | polygyne | 2 |
| BG240-W02 | BG240-W04 | 0.3682 | related | polygyne | 2 |
| BG240-W03 | BG240-W05 | 0.3778 | related | polygyne | 2 |
| BG240-W04 | BG240-W05 | 0.471 | related | polygyne | 2 |
| BG240-W02 | BG240-W05 | 0.3965 | related | polygyne | 2 |
| BG240-W01 | BG240-W05 | 0.388 | related | polygyne | 2 |
| BG240-W03 | BG240-W04 | 0.3691 | related | polygyne | 2 |
| BG240-W02 | BG240-W03 | 0.3912 | related | polygyne | 2 |
| BG240-W01 | BG240-W02 | 0.3967 | related | polygyne | 2 |
| BG241-W01 | BG241-W05 | 0.4099 | related | polygyne | 2 |
| BG241-W01 | BG241-W04 | 0.3908 | related | polygyne | 2 |
| BG241-W02 | BG241-W03 | 0.4093 | related | polygyne | 2 |
| BG241-W02 | BG241-W04 | 0.3815 | related | polygyne | 2 |
| BG241-W01 | BG241-W02 | 0.4015 | related | polygyne | 2 |
| BG241-W04 | BG241-W05 | 0.4052 | related | polygyne | 2 |
| BG241-W02 | BG241-W05 | 0.3997 | related | polygyne | 2 |
| BG241-W03 | BG241-W04 | 0.3766 | related | polygyne | 2 |
| BG241-W03 | BG241-W05 | 0.3892 | related | polygyne | 2 |
| BG241-W01 | BG241-W03 | 0.4269 | related | polygyne | 2 |
| BG270-W01 | BG270-W05 | 0.4974 | related | polygyne | 1 |
| BG270-W02 | BG270-W03 | 0.3752 | related | polygyne | 1 |
| BG270-W03 | BG270-W04 | 0.3374 | related | polygyne | 1 |
| BG270-W02 | BG270-W04 | 0.3489 | related | polygyne | 1 |
| BG270-W01 | BG270-W03 | 0.4394 | related | polygyne | 1 |
| BG270-W03 | BG270-W05 | 0.3359 | related | polygyne | 1 |
| BG270-W01 | BG270-W04 | 0.3931 | related | polygyne | 1 |
| BG270-W01 | BG270-W02 | 0.4787 | related | polygyne | 1 |
| BG270-W04 | BG270-W05 | 0.418 | related | polygyne | 1 |
| BG270-W02 | BG270-W05 | 0.4454 | related | polygyne | 1 |
| BG272-W03 | BG272-W05 | 0.4618 | related | polygyne | 1 |
| BG272-W01 | BG272-W03 | 0.3853 | related | polygyne | 1 |

| | | | | | |
|---|---|---|---|---|---|
| BG272-W01 | BG272-W05 | 0.3644 | related | polygyne | 1 |
| BG272-W02 | BG272-W03 | 0.4177 | related | polygyne | 1 |
| BG272-W01 | BG272-W02 | 0.3768 | related | polygyne | 1 |
| BG272-W02 | BG272-W05 | 0.442 | related | polygyne | 1 |
| BG233-W02 | BG233-W05 | 0.4415 | related | polygyne | 3 |
| BG233-W01 | BG233-W02 | 0.4015 | related | polygyne | 3 |
| BG233-W03 | BG233-W05 | 0.4408 | related | polygyne | 3 |
| BG233-W01 | BG233-W03 | 0.3934 | related | polygyne | 3 |
| BG233-W01 | BG233-W05 | 0.411 | related | polygyne | 3 |
| BG233-W03 | BG233-W04 | 0.3999 | related | polygyne | 3 |
| BG233-W04 | BG233-W05 | 0.4237 | related | polygyne | 3 |
| BG233-W02 | BG233-W03 | 0.4036 | related | polygyne | 3 |
| BG233-W02 | BG233-W04 | 0.4279 | related | polygyne | 3 |
| BG233-W01 | BG233-W04 | 0.3882 | related | polygyne | 3 |
| BG226-W03 | BG226-W04 | 0.4138 | related | polygyne | 3 |
| BG226-W02 | BG226-W05 | 0.4348 | related | polygyne | 3 |
| BG226-W02 | BG226-W04 | 0.4077 | related | polygyne | 3 |
| BG226-W02 | BG226-W03 | 0.4563 | related | polygyne | 3 |
| BG226-W01 | BG226-W03 | 0.4229 | related | polygyne | 3 |
| BG226-W01 | BG226-W04 | 0.3873 | related | polygyne | 3 |
| BG226-W04 | BG226-W05 | 0.4116 | related | polygyne | 3 |
| BG226-W03 | BG226-W05 | 0.396 | related | polygyne | 3 |
| BG226-W01 | BG226-W02 | 0.3929 | related | polygyne | 3 |
| BG226-W01 | BG226-W05 | 0.414 | related | polygyne | 3 |
| BG225-W02 | BG225-W04 | 0.4901 | related | polygyne | 3 |
| BG225-W03 | BG225-W05 | 0.3488 | related | polygyne | 3 |
| BG225-W03 | BG225-W04 | 0.4219 | related | polygyne | 3 |
| BG225-W02 | BG225-W03 | 0.4134 | related | polygyne | 3 |
| BG225-W02 | BG225-W05 | 0.4234 | related | polygyne | 3 |
| BG225-W01 | BG225-W03 | 0.4412 | related | polygyne | 3 |
| BG225-W01 | BG225-W02 | 0.4037 | related | polygyne | 3 |
| BG225-W04 | BG225-W05 | 0.4308 | related | polygyne | 3 |
| BG225-W01 | BG225-W04 | 0.4201 | related | polygyne | 3 |
| BG225-W01 | BG225-W05 | 0.4365 | related | polygyne | 3 |
| BG243-W02 | BG243-W04 | 0.5602 | full-sibs | polygyne | 2 |
| BG243-W04 | BG243-W05 | 0.3923 | related | polygyne | 2 |
| BG243-W02 | BG243-W03 | 0.5549 | full-sibs | polygyne | 2 |
| BG243-W01 | BG243-W02 | 0.3988 | related | polygyne | 2 |
| BG243-W01 | BG243-W03 | 0.4177 | related | polygyne | 2 |

| | | | | | |
|---|---|---|---|---|---|
| BG243-W03 | BG243-W04 | 0.545 | related | polygyne | 2 |
| BG243-W01 | BG243-W04 | 0.3803 | related | polygyne | 2 |
| BG243-W03 | BG243-W05 | 0.4627 | related | polygyne | 2 |
| BG243-W01 | BG243-W05 | 0.4009 | related | polygyne | 2 |
| BG243-W02 | BG243-W05 | 0.4022 | related | polygyne | 2 |
| BG237-W02 | BG237-W03 | 0.4262 | related | polygyne | 2 |
| BG237-W02 | BG237-W04 | 0.4408 | related | polygyne | 2 |
| BG237-W02 | BG237-W05 | 0.4578 | related | polygyne | 2 |
| BG237-W01 | BG237-W02 | 0.4345 | related | polygyne | 2 |
| BG237-W03 | BG237-W05 | 0.4967 | related | polygyne | 2 |
| BG237-W01 | BG237-W04 | 0.439 | related | polygyne | 2 |
| BG237-W03 | BG237-W04 | 0.4307 | related | polygyne | 2 |
| BG237-W04 | BG237-W05 | 0.4641 | related | polygyne | 2 |
| BG237-W01 | BG237-W03 | 0.4718 | related | polygyne | 2 |
| BG237-W01 | BG237-W05 | 0.4735 | related | polygyne | 2 |
| BG232-W03 | BG232-W04 | 0.4043 | related | polygyne | 3 |
| BG232-W03 | BG232-W05 | 0.3961 | related | polygyne | 3 |
| BG232-W01 | BG232-W05 | 0.4554 | related | polygyne | 3 |
| BG232-W02 | BG232-W04 | 0.5255 | related | polygyne | 3 |
| BG232-W02 | BG232-W03 | 0.4551 | related | polygyne | 3 |
| BG232-W01 | BG232-W04 | 0.4838 | related | polygyne | 3 |
| BG232-W02 | BG232-W05 | 0.5201 | related | polygyne | 3 |
| BG232-W01 | BG232-W03 | 0.5264 | related | polygyne | 3 |
| BG232-W04 | BG232-W05 | 0.612 | full-sibs | polygyne | 3 |
| BG232-W01 | BG232-W02 | 0.4532 | related | polygyne | 3 |
| BG242-W01 | BG242-W03 | 0.5144 | related | polygyne | 2 |
| BG242-W01 | BG242-W04 | 0.5166 | related | polygyne | 2 |
| BG242-W03 | BG242-W04 | 0.7009 | full-sibs | polygyne | 2 |
| BG242-W04 | BG242-W05 | 0.6066 | full-sibs | polygyne | 2 |
| BG242-W03 | BG242-W05 | 0.5452 | related | polygyne | 2 |
| BG242-W01 | BG242-W05 | 0.5251 | related | polygyne | 2 |

**Table S3.** Cross-validation (CV) error estimates ($K$ = 1-10) for the *F. paralugubris* ADMIXTURE analysis. CV error, as determined by ADMIXTURE, suggests the best number of ancestries is $K$ = 3 (in bold).

| *K* | Cross-validation error |
|---|---|
| 1 | 0.40212 |
| 2 | 0.38212 |

| | |
|---|---|
| **3** | **0.37798** |
| 4 | 0.37870 |
| 5 | 0.38700 |
| 6 | 0.39017 |
| 7 | 0.39815 |
| 8 | 0.41252 |
| 9 | 0.42154 |
| 10 | 0.43617 |

**Table S4.** Intra-nest relatedness as estimated by *COANCESTRY* using the Wang estimator in *F. aquilonia*. These results are visualized in the main text in Figure 2B.

| **Ind1** | **Ind2** | **Wang estimator** | **Parentage** | **Nest social form** |
|---|---|---|---|---|
| BG229.95-W01 | BG229.95-W02 | 0.361 | related | polygyne |
| BG229.95-W01 | BG229.95-W03 | 0.4868 | related | polygyne |
| BG229.95-W01 | BG229.95-W04 | 0.3559 | related | polygyne |
| BG229.95-W01 | BG229.95-W05 | 0.5314 | related | polygyne |
| BG229.95-W01 | BG229.95-W1 | 0.2308 | related | polygyne |
| BG229.95-W01 | BG229.95-W2 | 0.1356 | not-sibs | polygyne |
| BG229.95-W01 | BG229.95-W3 | 0.2811 | related | polygyne |
| BG229.95-W02 | BG229.95-W03 | 0.3997 | related | polygyne |
| BG229.95-W02 | BG229.95-W04 | 0.3626 | related | polygyne |
| BG229.95-W02 | BG229.95-W05 | 0.3585 | related | polygyne |
| BG229.95-W02 | BG229.95-W1 | 0.1591 | not-sibs | polygyne |
| BG229.95-W02 | BG229.95-W2 | 0.1779 | not-sibs | polygyne |
| BG229.95-W02 | BG229.95-W3 | 0.2131 | related | polygyne |
| BG229.95-W03 | BG229.95-W04 | 0.3153 | related | polygyne |
| BG229.95-W03 | BG229.95-W05 | 0.4808 | related | polygyne |
| BG229.95-W03 | BG229.95-W1 | 0.1259 | not-sibs | polygyne |
| BG229.95-W03 | BG229.95-W2 | 0.1044 | not-sibs | polygyne |
| BG229.95-W03 | BG229.95-W3 | 0.4878 | related | polygyne |
| BG229.95-W04 | BG229.95-W05 | 0.3159 | related | polygyne |
| BG229.95-W04 | BG229.95-W1 | 0.1343 | not-sibs | polygyne |
| BG229.95-W04 | BG229.95-W2 | 0.1251 | not-sibs | polygyne |
| BG229.95-W04 | BG229.95-W3 | 0.1281 | not-sibs | polygyne |
| BG229.95-W05 | BG229.95-W1 | 0.1998 | related | polygyne |
| BG229.95-W05 | BG229.95-W2 | 0.1512 | not-sibs | polygyne |
| BG229.95-W05 | BG229.95-W3 | 0.2649 | related | polygyne |
| BG229.95-W1 | BG229.95-W2 | 0.3915 | related | polygyne |
| BG229.95-W1 | BG229.95-W3 | 0.4077 | related | polygyne |
| BG229.95-W2 | BG229.95-W3 | 0.3749 | related | polygyne |

| | | | | |
|---|---|---|---|---|
| BG231-W01 | BG231-W02 | 0.42 | related | monogyne, polyandrous |
| BG231-W01 | BG231-W03 | 0.4495 | related | monogyne, polyandrous |
| BG231-W01 | BG231-W04 | 0.7027 | full-sibs | monogyne, polyandrous |
| BG231-W01 | BG231-W05 | 0.4623 | related | monogyne, polyandrous |
| BG231-W02 | BG231-W03 | 0.694 | full-sibs | monogyne, polyandrous |
| BG231-W02 | BG231-W04 | 0.401 | related | monogyne, polyandrous |
| BG231-W02 | BG231-W05 | 0.4265 | related | monogyne, polyandrous |
| BG231-W03 | BG231-W04 | 0.4115 | related | monogyne, polyandrous |
| BG231-W03 | BG231-W05 | 0.4525 | related | monogyne, polyandrous |
| BG231-W04 | BG231-W05 | 0.4083 | related | monogyne, polyandrous |
| BG247-W01 | BG247-W02 | 0.4249 | related | monogyne, polyandrous |
| BG247-W01 | BG247-W03 | 0.4381 | related | monogyne, polyandrous |
| BG247-W01 | BG247-W04 | 0.4141 | related | monogyne, polyandrous |
| BG247-W01 | BG247-W05 | 0.429 | related | monogyne, polyandrous |
| BG247-W02 | BG247-W03 | 0.6546 | full-sibs | monogyne, polyandrous |
| BG247-W02 | BG247-W04 | 0.6665 | full-sibs | monogyne, polyandrous |
| BG247-W02 | BG247-W05 | 0.681 | full-sibs | monogyne, polyandrous |
| BG247-W03 | BG247-W04 | 0.664 | full-sibs | monogyne, polyandrous |
| BG247-W03 | BG247-W05 | 0.6644 | full-sibs | monogyne, polyandrous |
| BG247-W04 | BG247-W05 | 0.6753 | full-sibs | monogyne, polyandrous |
| BG245-W01 | BG245-W02 | 0.6426 | full-sibs | monogyne, monandrous |
| BG245-W01 | BG245-W03 | 0.615 | full-sibs | monogyne, monandrous |
| BG245-W01 | BG245-W04 | 0.6364 | full-sibs | monogyne, monandrous |
| BG245-W01 | BG245-W05 | 0.6448 | full-sibs | monogyne, monandrous |
| BG245-W02 | BG245-W03 | 0.7053 | full-sibs | monogyne, monandrous |
| BG245-W02 | BG245-W04 | 0.6923 | full-sibs | monogyne, monandrous |
| BG245-W02 | BG245-W05 | 0.6875 | full-sibs | monogyne, monandrous |
| BG245-W03 | BG245-W04 | 0.7035 | full-sibs | monogyne, monandrous |
| BG245-W03 | BG245-W05 | 0.6992 | full-sibs | monogyne, monandrous |
| BG245-W04 | BG245-W05 | 0.6707 | full-sibs | monogyne, monandrous |

**Table S5.** Intra-nest relatedness as estimated by *COANCESTRY* using the Wang estimator in *F. truncorum*. These results are visualized in the main text in Figure 2C.

| Ind1 | Ind2 | Wang estimator | Parentage | Nest social form |
|---|---|---|---|---|
| BG227-W01 | BG227-W02 | 0.6727 | full-sibs | monogyne, monandrous |
| BG227-W01 | BG227-W03 | 0.6495 | full-sibs | monogyne, monandrous |
| BG227-W01 | BG227-W04 | 0.6607 | full-sibs | monogyne, monandrous |
| BG227-W01 | BG227-W05 | 0.6894 | full-sibs | monogyne, monandrous |

| | | | | |
|---|---|---|---|---|
| BG227-W02 | BG227-W03 | 0.6593 | full-sibs | monogyne, monandrous |
| BG227-W02 | BG227-W04 | 0.6457 | full-sibs | monogyne, monandrous |
| BG227-W02 | BG227-W05 | 0.6636 | full-sibs | monogyne, monandrous |
| BG227-W03 | BG227-W04 | 0.6725 | full-sibs | monogyne, monandrous |
| BG227-W03 | BG227-W05 | 0.6646 | full-sibs | monogyne, monandrous |
| BG227-W04 | BG227-W05 | 0.6399 | full-sibs | monogyne, monandrous |
| BG44-W1 | BG44-W2 | 0.7066 | full-sibs | monogyne, monandrous |
| BG44-W1 | BG44-W3 | 0.6253 | full-sibs | monogyne, monandrous |
| BG44-W1 | BG44-W4 | 0.7325 | full-sibs | monogyne, monandrous |
| BG44-W2 | BG44-W3 | 0.5748 | full-sibs | monogyne, monandrous |
| BG44-W2 | BG44-W4 | 0.6888 | full-sibs | monogyne, monandrous |
| BG44-W3 | BG44-W4 | 0.6658 | full-sibs | monogyne, monandrous |
| BG81-W1 | BG81-W2 | 0.7139 | full-sibs | monogyne, monandrous |
| BG81-W1 | BG81-W3 | 0.7081 | full-sibs | monogyne, monandrous |
| BG81-W1 | BG81-W4 | 0.774 | full-sibs | monogyne, monandrous |
| BG81-W2 | BG81-W3 | 0.6908 | full-sibs | monogyne, monandrous |
| BG81-W2 | BG81-W4 | 0.6943 | full-sibs | monogyne, monandrous |
| BG81-W3 | BG81-W4 | 0.7469 | full-sibs | monogyne, monandrous |
